# Membrane-type 1 matrix metalloproteinase (MMP-14) modulates tissue homeostasis by a non-proteolytic mechanism

**DOI:** 10.1101/631739

**Authors:** Mukundan Attur, Cuijie Lu, Xiaodong Zhang, Tianzhen Han, Cassidy Alexandre, Cristina Valacca, Shuai Zheng, Sarina Meikle, Branka Brukner Dabovic, Evelyne Tassone, Qing Yang, Victoria Kolupaeva, Shoshana Yakar, Steven Abramson, Paolo Mignatti

**Affiliations:** Department of Medicine, Division of Rheumatology, NYU School of Medicine, 550 First Avenue, New York, NY 10016, USA; Department of Cardiothoracic Surgery, NYU School of Medicine, 550 First Avenue, New York, NY 10016, USA; Department of Cell Biology, NYU School of Medicine, 550 First Avenue, New York, NY 10016, USA; Department of Microbiology, NYU School of Medicine, 550 First Avenue, New York, NY 10016, USA; Department of Basic Science & Craniofacial Biology, NYU College of Dentistry, 345 E. 24th Street, New York, NY 10010, USA

## Abstract

Membrane-type 1 matrix metalloproteinase (MT1-MMP, MMP-14), a transmembrane proteinase with a short cytoplasmic tail, is a major effector of extracellular matrix (ECM) remodeling. Genetic silencing of MT1-MMP in mouse (*Mmp14^−/−^*) and man causes dwarfism, osteopenia, arthritis and lipodystrophy, abnormalities ascribed to defective collagen turnover. We have previously shown non-proteolytic functions of MT1-MMP mediated by its cytoplasmic tail, where the unique tyrosine (Y573) controls intracellular signaling. The Y573D mutation blocks TIMP2/MT1-MMP-induced Erk1/2 and Akt signaling without affecting proteolytic activity. Here we report that a mouse with the MT1-MMP Y573D mutation (*Mmp14^Y573D/Y573D^*) shows abnormalities similar to, but also different from those of *Mmp14^−/−^* mice. Skeletal stem cells (SSC) of *Mmp14^Y573D/Y573D^* mice show defective differentiation consistent with the mouse phenotype, which is rescued by wild-type SSC transplant. These results provide the first *in vivo* demonstration that MT1-MMP modulates bone, cartilage and fat homeostasis by controlling SSC differentiation through a mechanism independent of proteolysis.

## INTRODUCTION

Membrane-type 1 matrix metalloproteinase (MT1-MMP, or MMP-14), the product of the gene *MMP14*, is a cell-membrane-bound proteinase with an extracellular catalytic site and a 20-amino acid cytoplasmic tail (Itoh, 2015; Sato et al., 1994). It degrades a variety of ECM components and is expressed by a wide array of normal and tumor cells. Notably, MT1-MMP is the only MMP whose genetic deficiency in the mouse results in severe phenotypes and early death. These features have implicated MT1-MMP as an important component of the proteolytic mechanisms of physiological and pathological processes including bone, cartilage and adipose tissue homeostasis, as well as tumor invasion, angiogenesis and metastasis. The analysis of the phenotype of mice genetically deficient in MT1-MMP (*Mmp14^−/−^*) has shown key roles of this proteinase in the postnatal development and growth of cartilage, bone and adipose tissue. *Mmp14* deficiency in the mouse results in dwarfism, severe osteopenia, generalized arthritis and lipodystrophy, among other abnormalities (Chun et al., 2006; Chun & Inoue, 2014; Holmbeck et al., 1999; Zhou et al., 2000). In humans a mutation of *MMP14* causes multicentric osteolysis and arthritis disease, or Winchester syndrome, which recapitulates much of the phenotype of the *Mmp14^−/−^* mouse (Evans et al., 2012). Conditional *Mmp14* knockout in uncommitted skeletal stem cells (SSC, also referred to as mesenchymal stem cells), the common progenitors of osteoblasts, chondrocytes and adipocytes, recapitulates the skeletal phenotype of the global *Mmp14* knockout mouse (Y. Tang et al., 2013). Conversely, unlike global *Mmp14* deficiency conditional *Mmp14* knockout in SSC results in thickening of articular cartilage and increased bone marrow (BM)-associated fat but decreased subcutaneous fat, which derives from distinct progenitor cells (Chun et al., 2006; Gupta et al., 2012; Tran et al., 2012). In light of the fundamental role of MT1-MMP in ECM degradation, it has been proposed that the phenotypes of *Mmp14^−/−^* mice result from defective collagen turnover (Chun et al., 2006; Chun & Inoue, 2014; Holmbeck et al., 1999; Zhou et al., 2000).

However, a number of *in vitro* studies have provided evidence for a variety of non-proteolytic roles of MT1-MMP. We have previously shown that the MT1-MMP cytoplasmic tail activates Ras-ERK1/2 and Ras-AKT signaling by a non-proteolytic mechanism that controls cell proliferation, migration and apoptosis *in vitro*, as well as tumor growth *in vivo* (D’Alessio et al., 2008; Valacca et al., 2015). Signaling is activated in a dose- and time-dependent manner by MT1-MMP binding of low nanomolar concentrations of tissue inhibitor of metalloproteinases-2 (TIMP-2), a physiological protein inhibitor of MT1-MMP. Signaling activation is also mediated by mutant TIMP-2 lacking MMP inhibitory activity (Ala+ TIMP-2) as well as by mutant MT1-MMP devoid of proteolytic activity (MT1-MMP E240A), showing that the signaling mechanism is proteolysis-independent. We also showed that MT1-MMP signaling requires the unique tyrosine (Y573) in the cytoplasmic tail, which is phosphorylated by Src and LIM1 kinases (Lagoutte et al., 2016; Nyalendo et al., 2007), and that Y573 substitution with aspartic acid (Y573D), a negatively charged amino acid like phosphotyrosine, abrogates MT1-MMP-mediated activation of Ras-ERK1/2 (D’Alessio et al., 2008). Y573 is required for activation of the small GTPase Rac1 (Gonzalo et al., 2010), controls macrophage migration and infiltration at sites of inflammation (Gonzalo et al., 2010; Sakamoto & Seiki, 2009), as well as MT1-MMP interaction with Src and focal adhesion kinase (FAK) (Y. Wang & McNiven, 2012).

Based on these observations, we generated a mutant mouse with the Y573D substitution in the cytoplasmic tail of MT1-MMP (MT1-MMP Y573D). Here we report that this mouse shows abnormalities in postnatal bone, cartilage and adipose tissue development and growth that partly recapitulate the phenotype of the *MMP14^−/−^* mouse, but also significant dissimilarities that indicate the relevance of the balance between MT1-MMP proteolytic activity and proteolysis-independent signaling. These phenotypes derive from dysregulation of BM SSC differentiation, and are rescued by wild-type (wt) BM transplant.

## RESULTS

### Generation and macroscopic characterization of MT1-MMP Y573D mice

Mice heterozygous (*Mmp^Y573D/wt^*) or homozygous (*Mmp^Y573D/Y573D^*) for the MT1-MMP Y573D mutation, generated as described in Transparent Methods (Fig. 1 A and B), were macroscopically indistinguishable from *Mmp14^wt/wt^* mice at birth. However, after the first 2 months *Mmp14^Y573D/wt^* and *Mmp14^Y573D/Y573D^* mice showed a slightly lower growth rate than their *Mmp14^wt/wt^* littermates (Fig. 1 C). In mice older than 3 months the weight of *Mmp14^Y573D/Y573D^* male and female mice was 15 % lower than that of age- and sex-matched *Mmp14^wt/wt^* littermates (Fig. 1 D). Adult *Mmp14^Y573D/Y573D^* mice showed no macroscopic morphological alterations. Both male and female *Mmp14^Y573D/Y573D^* mice were fertile, bred and lived up to > 2 years of age similarly to *Mmp14^wt/wt^* mice. Thus, the macroscopic phenotype of *Mmp14^wt/wt^* mice did not show the dramatic abnormalities of *Mmp14^−/−^* mice, which are up to 25% smaller than their wt littermate, have striking skeletal defects and a shortened life span (Table 1).

**Figure 1.**
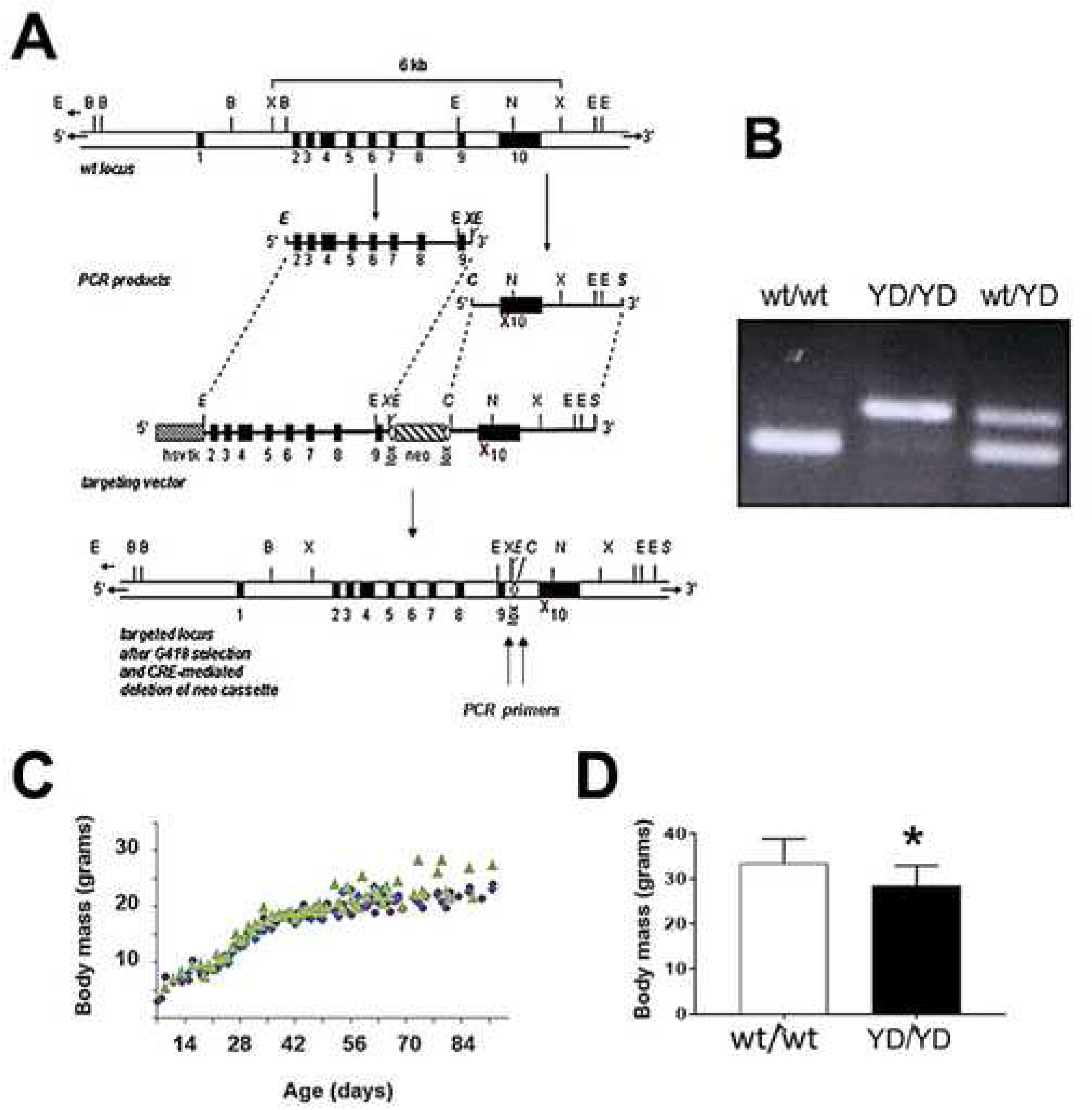
Generation and macroscopic characterization of the MT1-MMP Y573D mouse. **A.** Schematic representation of the targeting vector and targeted locus. To construct the targeting vector the genomic sequence of mouse MT1-MMP containing exons 2 to 10 was PCR amplified from W4 embryonic stem (ES) cells’ DNA. A floxed neomycin resistance cassette (neo) was inserted into intron 9, adjacent to the 5’ end of exon 10, which encodes Y573, and a *hsv*-thymidine kinase cassette (hsvtk) was placed at the 5′ end of the construct. The TAC codon for Y573 was mutated into GAC by PCR. The construct was electroporated into W4 ES cells and colonies of recombinant cells were selected as described in Transparent Methods. The red X indicates the Y573D substitution; B: BamHI; C: ClaI; E: EcoRI; N: NotI; S: SalI; X: XhoI. **B.** PCR genotyping. The mice were genotyped by PCR using primers flanking the *loxP* sites, which afford identification of the three genotypes. The sequence of the primers is reported in Transparent Methods. **C.** Growth curve of *Mmp14^wt/wt^* (▲), *Mmp14^wt/Y573D^* (▼) and (*Mmp14^Y573D/Y573D^* (●) littermates. **D.** Body mass of 5- to 7-month old *Mmp14^wt/wt^* (wt/wt; n = 14) and *Mmp14^Y573D/Y573D^* (YD/YD; n = 15) mice of both sexes (33.54 ± 1.44 g vs. 28.5 ± 1.15 g, respectively). *: p ≤ 0.001.

**Table 1.** Comparison of the phenotypes of MT1-MMP mutant mice. **1.** The Cartoon mouse harbors a S466P point mutation in the MT1-MMP hemopexin domain that generates a misfolded, temperature-sensitive mutant that retained in the endoplasmic reticulum (Sakr et al., 2018).

| **Genotype** | **Global knockout** | **SSC Dermo-1** | **Cartoon<sup>1</sup>** | **Y573D** |
| --- | --- | --- | --- | --- |
| **Life span** | 2-3 months | 4-5 months | 3-6 weeks | Normal (2 years) |
| **Generalized fibrosis of soft tissues** | +++ | Unknown | Unknown | - |
| **Skeleton** | Dwarfism<br>Craniofacial abnormalities: domed skull and short snout<br>Osteopenia | Dwarfism<br>Craniofacial abnormalities: domed skull and short snout<br>Osteopenia | Dwarfism<br>Craniofacial abnormalities: domed skull and short snout<br>Osteopenia | Decreased cortical bone but increased trabecular bone |
| **Joints** | Generalized arthritis with degeneration of articular cartilage | Increased cartilage thickness | Increased cartilage thickness | Decreased cartilage thickness<br>Osteoarthritis-like lesions |
| **Adipose tissue** | Generalized lipodystrophy<br>Normal brown fat | Increased bone marrow fat<br>Decreased sub-cutaneous fat<br>Brown fat unknown | Generalized lipodystrophy<br>Brown fat unknown | Generalized lipodystrophy<br>Brown fat hypertrophy |
| **Inflammation** | Increase in inflammatory cytokines | Unknown | Unknown | Decrease of Inflammatory cytokines |
| **Hematopoiesis** | Pancytopenia | Unknown | Unknown | Normal levels of circulating blood cells |
| **SSC differentiation in 2D culture** | Unknown | Decreased osteogenesis<br>Increased adipogenesis and chondrogenesis | Decreased osteogenesis<br>Increased adipogenesis (both in 3D collagen culture) | Increased osteogenesis<br>Decreased adipogenesis and chondrogenesis |

**Table 1. Comparison of the phenotypes of MT1-MMP mutant mice.**
| **Genotype** | **Global knockout** | **SSC Dermo-1** | **Cartoon<sup>1</sup>** | **Y573D** |
| --- | --- | --- | --- | --- |
| **Life span** | 2-3 months | 4-5 months | 3-6 weeks | Normal (2 years) |
| **Generalized fibrosis of soft tissues** | +++ | Unknown | Unknown | - |
| **Skeleton** | Dwarfism<br>Craniofacial abnormalities:<br>domed skull and short snout<br>Osteopenia | Dwarfism<br>Craniofacial abnormalities:<br>domed skull and short snout<br>Osteopenia | Dwarfism<br>Craniofacial abnormalities:<br>domed skull and short snout<br>Osteopenia | Decreased cortical bone but increased trabecular bone |
| **Joints** | Generalized arthritis with degeneration of articular cartilage | Increased cartilage thickness | Increased cartilage thickness | Decreased cartilage thickness<br>Osteoarthritis-like lesions |
| **Adipose tissue** | Generalized lipodystrophy<br>Normal brown fat | Increased bone marrow fat<br>Decreased sub-cutaneous fat<br>Brown fat unknown | Generalized lipodystrophy<br>Brown fat unknown | Generalized lipodystrophy<br>Brown fat hypertrophy |
| **Inflammation** | Increase in inflammatory cytokines | Unknown | Unknown | Decrease of Inflammatory cytokines |
| **Hematopoiesis** | Pancytopenia | Unknown | Unknown | Normal levels of circulating blood cells |
| **SSC differentiation in 2D culture** | Unknown | Decreased osteogenesis<br>Increased adipogenesis and chondrogenesis | Decreased osteogenesis<br>Increased adipogenesis (both in 3D collagen culture) | Increased osteogenesis<br>Decreased adipogenesis and chondrogenesis |

### The Y573D mutation does not affect the proteolytic activity of MT1-MMP

A major consequence of the genetic deficiency of MT1-MMP collagenolytic activity in *Mmp14^−/−^* mice is the progressive development of fibrosis of soft tissues, including fibrosis of the dermis and hair follicles, with subsequent hair loss (Holmbeck et al, 1999). *Mmp14^Y573D/Y573D^* mice up to 2 years of age showed no hair loss, and histological analysis of their skin showed no fibrosis of the dermis or hair follicles (Fig. 2 A), indicating that the collagenolytic activity of MT1-MMP Y573D *in vivo* is comparable to that of wt MT1-MMP. Consistent with this finding, primary fibroblasts from 3-weeks old *Mmp14^Y573D/Y573D^* and *Mmp14^wt/wt^* mice, which had comparable levels of MT1-MMP (Fig. 2 B) showed a similar capacity to degrade collagen, invade 3D collagen and activate proMMP-2 (Fig. 2 C – G). Because mutations of the cytoplasmic tail can affect MT1-MMP sorting to, and recycling from the cell membrane (Uekita et al., 2001), we also characterized the cell surface levels of MT1-MMP *in vivo* by flow cytometric analysis of circulating white blood monocytes from *Mmp14^Y573D/Y573D^* and *Mmp14^wt/wt^* mice. Consistent with our analysis of the cell-associated collagenolytic activity *in vitro*, the results (Figure S1 A) showed that cells of both genotypes had comparable levels of cell membrane-associated MT1-MMP, indicating that the Y573D mutation has minimal or no detectable effect on cell membrane expression of MT1-MMP *in vivo*.

**Figure 2.**
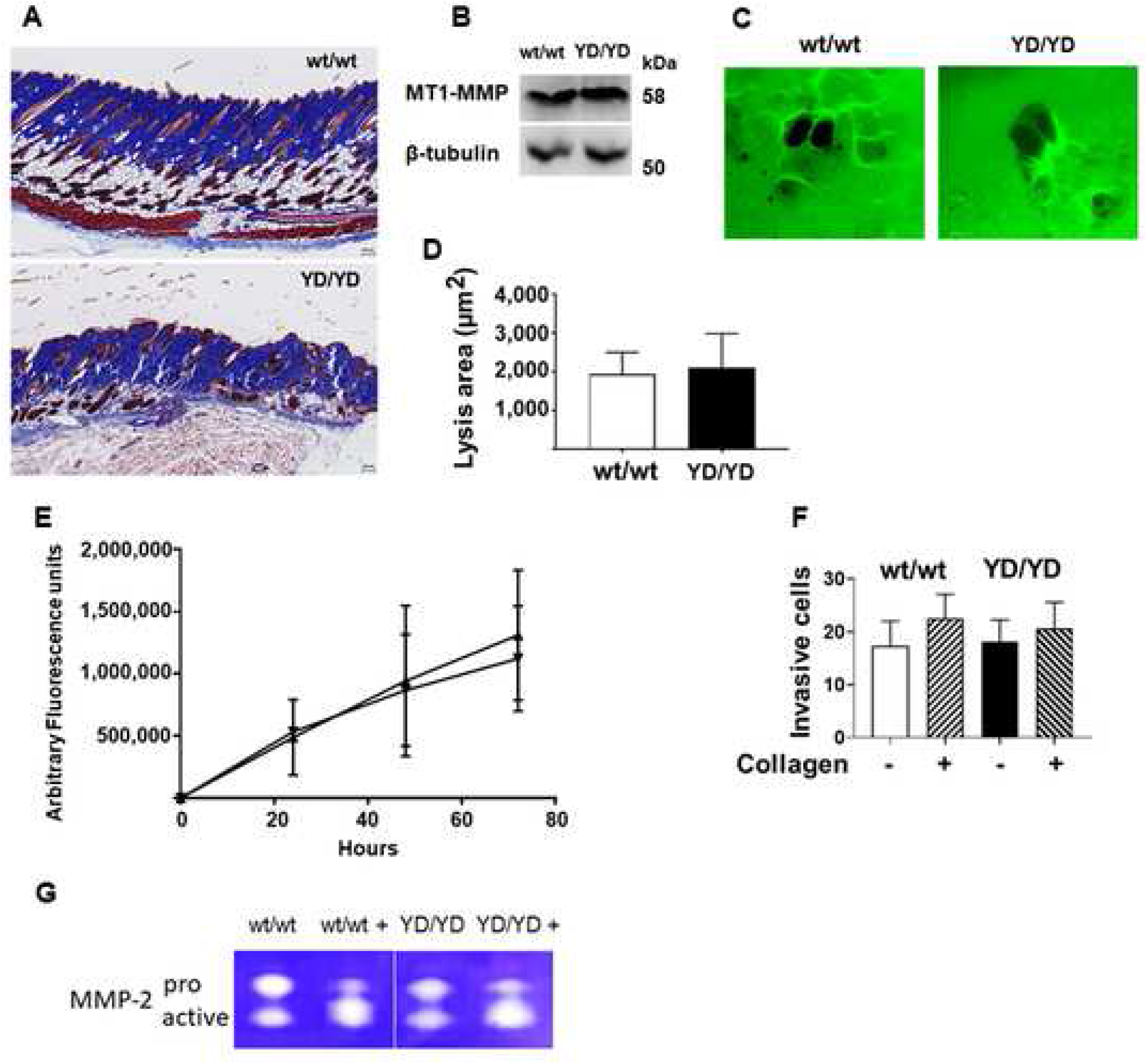
The Y573D mutation does not affect the proteolytic activity of MT1-MMP. **A.** Representative histological sections of the skin of 3-months old Mmp14wt/wt (wt/wt; top panel) and Mmp14Y573D/Y573D mice (YD/YD; bottom panel). H&E staining; original magnification: 4X. **B.** Western blotting analysis of MT1-MMP expression in primary fibroblasts from Mmp14wt/wt (wt/wt) and Mmp14Y573D/Y573D mice (YD/YD). β-tubulin is shown in the lower panel as a loading control. **C.** Representative images of primary fibroblasts from Mmp14wt/wt (wt/wt) and Mmp14Y573D/Y573D mice (YD/YD) grown atop Alexa Fluor 488-labeled collagen fibrils for 72 h. Areas of collagen degradation appear as black holes in the fluorescent collagen layer. Original magnification: 20 X. **D.** Lysis areas (wt/wt n = 60; YD/YD n = 76) were measured by Photoshop. Mean ± standard deviation is shown. This experiment was repeated three times with comparable results. **E.** Confluent primary fibroblasts from Mmp14wt/wt (▲) and Mmp14Y573D/ Y573D mice (▼) were grown atop Alexa Fluor 488-labeled collagen fibrils for 72 h, and the fluorescence released into the culture supernatant was measured at the indicated times as described in Methods. Mean and standard deviation of two independent experiments are shown. **F.** Analysis of cell migration and collagen invasion by primary fibroblasts from Mmp14^wt/wt^ (wt/wt) and ^Mmp14Y573D/Y573D^ mice (YD/YD) on 8-μm pore membranes coated (+) or non-coated (-) with type I collagen as described in Transparent Methods. Shown in the ordinate is the mean ± s.d. of the number of migrated cells per 20X microscopic field. This experiment was repeated twice with comparable results. **G.** Gelatin zymographic analysis of proMMP-2 activation in the serum-free conditioned medium of primary cultures of Mmp14wt/wt (wt/wt) and Mmp14Y573D/Y573D (YD/YD) mouse fibroblasts incubated with exogenous human recombinant proMMP-2 for 16 h. Before zymography the conditioned medium was incubated in the absence or presence (+) of APMA (1 mM in DMSO) for 1.5 h at 37° C as a positive control for proMMP-2 activation. This experiment was repeated three times with comparable results. See also Figure S1.

In addition to degrading collagen and other ECM components, MT1-MMP cleaves a variety of cell membrane proteins including the collagen receptors CD44 and discoidin domain receptor 1 (DDR1), as well as the metalloproteinase ADAM9 (Chan et al., 2012; Fu et al., 2013; Kajita et al., 2001). Fat, cartilage and bone (Fig. 3 A), the tissues affected by the MT1-MMP Y573D mutation (as described under the following subheading), as well as other tissues (Figure S1 B), showed minor variations in the levels of MT1-MMP expression in mice of the two genotypes. In fat MT1-MMP was expressed almost exclusively in the proenzyme form (62 kDa), whereas in cartilage and bone the 42-44 kDa autocatalytic degradation product was prevalent, indicating high levels of proteolytic activity (Rozanov et al., 2001).

**Figure 3.**
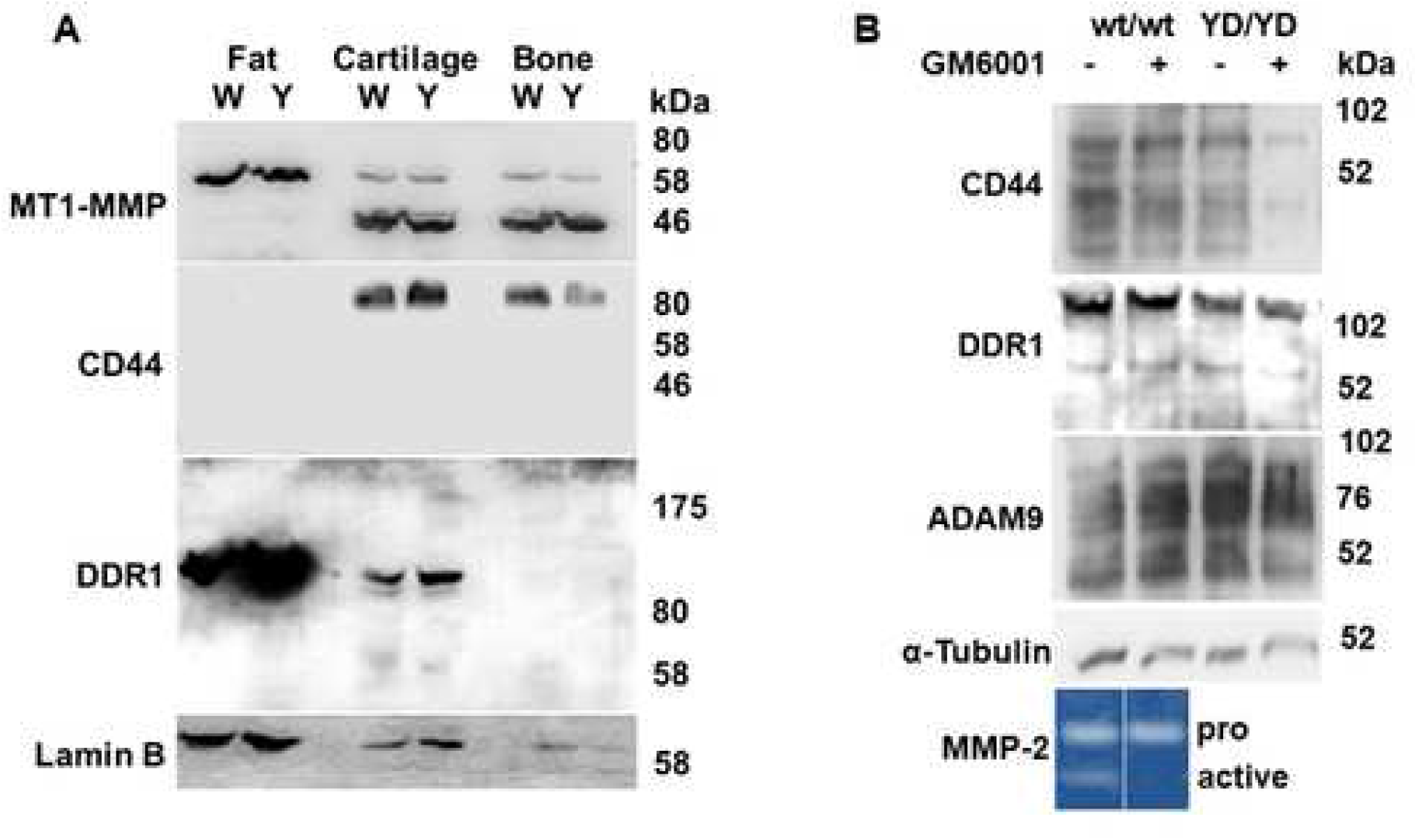
The Y573D mutation does not affect MT1-MMP cleavage of CD44, DDR1 or ADAM9. **A**. Western blotting analysis of MT1-MMP, CD44 and DDR1 expression and degradation in abdominal fat, joint cartilage and bone (femur) from 3-month old *Mmp14^wt/wt^* (W) and *Mmp14^Y573D/Y573D^* (Y) male mice. Tissues from three littermates per genotype were pooled for protein extraction. Lamin B is shown as a loading control. A representative result of multiple experiments is shown. **B.** Western blotting analysis of CD44, DDR1 and ADAM9 in cell extracts of primary fibroblasts from *Mmp14^wt/wt^* (wt/wt) and *Mmp14^Y573D/Y573D^* (YD/YD) mice grown in the presence (+) or absence (-) of GM6001 (50 μM) for 24 h. α-tubulin is shown as a loading control. A representative result of multiple experiments is shown. Bottom panel: gelatin zymography analysis of proMMP-2 activation is shown as a control for GM6001 inhibition of MT1-MMP. The cells were grown for 16 h in serum-free medium supplemented with human recombinant proMMP-2 without (-) or with (+) GM6001 (50 μM), and the conditioned medium was analyzed by gelatin zymography. See also Figure S1.

We could not detect CD44 in the fat of our mice (Fig. 3 A), and qPCR analysis showed extremely low levels of CD44 mRNA in both *Mmp14^Y573D/Y573D^* and *Mmp14^wt/wt^* mice (Ct values 32.1 and 37.9, respectively). However, the 80 kDa native form of CD44 was present in the cartilage and bone of *Mmp14^Y573D/Y573D^* and *Mmp14^wt/wt^* mice in comparable amounts, and no degradation products were detected in mice of either genotype. DDR1 could not be detected in the bone of *Mmp14^Y573D/Y573D^* or *Mmp14^wt/wt^* mice (Fig. 3 A), a result consistent with qPCR analysis (Ct values 31.89 and 31.76, respectively). Conversely, DDR1 was detected in fat and cartilage, but not in bone, extracts as a ~ 120 kDa form. Comparable levels of a ~ 62 - 65 kDa form consistent with a degradation product resulting from MT1-MMP proteolysis (Fu et al., 2013) were present in cartilage, whereas higher levels of ~ 120 kDa DDR1 was expressed in *Mmp14^Y573D/Y573D^* than in *Mmp14^wt/wt^* fat. No other degradation products were detected, showing that wt MT1-MMP and MT1-MMP Y573D have comparable capacity to cleave CD44 and DDR1 *in vivo*.

To corroborate these findings we analyzed CD44, DDR1 and ADAM9 in primary fibroblasts from adult *Mmp14^Y573D/Y573D^* and *Mmp14^wt/wt^* mice (Fig. 3 B). Western blotting analysis of cell extracts showed comparable levels of native, 80 kDa CD44 and lower molecular weight immunoreactive bands. *Mmp14^Y573D/Y573D^* and *Mmp14^wt/wt^* cell extracts showed similar levels of native, ~ 120 kDa DDR1 and ~ 62 – 65 kDa bands consistent with the degradation product resulting from MT1-MMP proteolysis (Fu et al., 2013), as well as ~ 90 kDa ADAM9 (Fig. 3 B) and ~ 40 – 60 kDa immunoreactive bands consistent with ADAM-9 degradation products generated by MT1-MMP (Chan, et al., 2012). However, similar CD44 and DDR1 bands were observed in extracts of cells grown in the presence or absence of the MMP inhibitor GM6001 (Ilomastat), indicating that they either represent nonspecific bands or degradation products generated by non-MMP mediated cleavage. Therefore, these results showed that the Y573D mutation does not affect MT1-MMP cleavage of CD44, DDR1 or ADAM9 *in vitro* or *in vivo*.

### MT1-MMP Y573D mice show structural and gene expression abnormalities in bone, articular cartilage and adipose tissue

A striking phenotype of *Mmp14^−/−^* mice is the dramatically decreased length of long bones, with severe osteopenia and arthritis (Holmbeck et al., 1999; Zhou et al., 2000). At 5 months of age the femurs of *Mmp14^Y573D/Y573D^* mice were only ~ 5% shorter than those of *Mmp14^wt/wt^* mice (14.81 ± 0.1519 mm *vs.* 15.61 ± 0.0619 mm; n=10; p = 0.0001). However, cortical bone thickness at the femur mid-diaphysis was markedly decreased (20-25%) in *Mmp14^Y573D/Y573D^ vs. Mmp14^wt/wt^* mice (Fig. 4 A), a finding consistent with the osteopenia of the *Mmp14^−/−^* mouse (Holmbeck et al., 1999; Zhou et al., 2000). Conversely, unlike *Mmp14^−/−^* mice, which have reduced trabecular bone, and in the homozygous but not in the heterozygous state (Holmbeck et al., 1999; Zhou et al., 2000), femurs and tibias showed significantly increased (40-50%) trabecular bone in both *Mmp14^Y573D/Y573D^* and *Mmp14^Y573D/wt^* mice relative to age- and sex-matched *Mmp14^wt/wt^* mice (Fig. 4 B - D). This effect was accompanied by increased TRAP-positive osteoclasts (Figure S2), indicating enhanced bone remodeling, a feature also observed in *Mmp14^−/−^* mice (Holmbeck et al., 1999).

**Figure 4.**
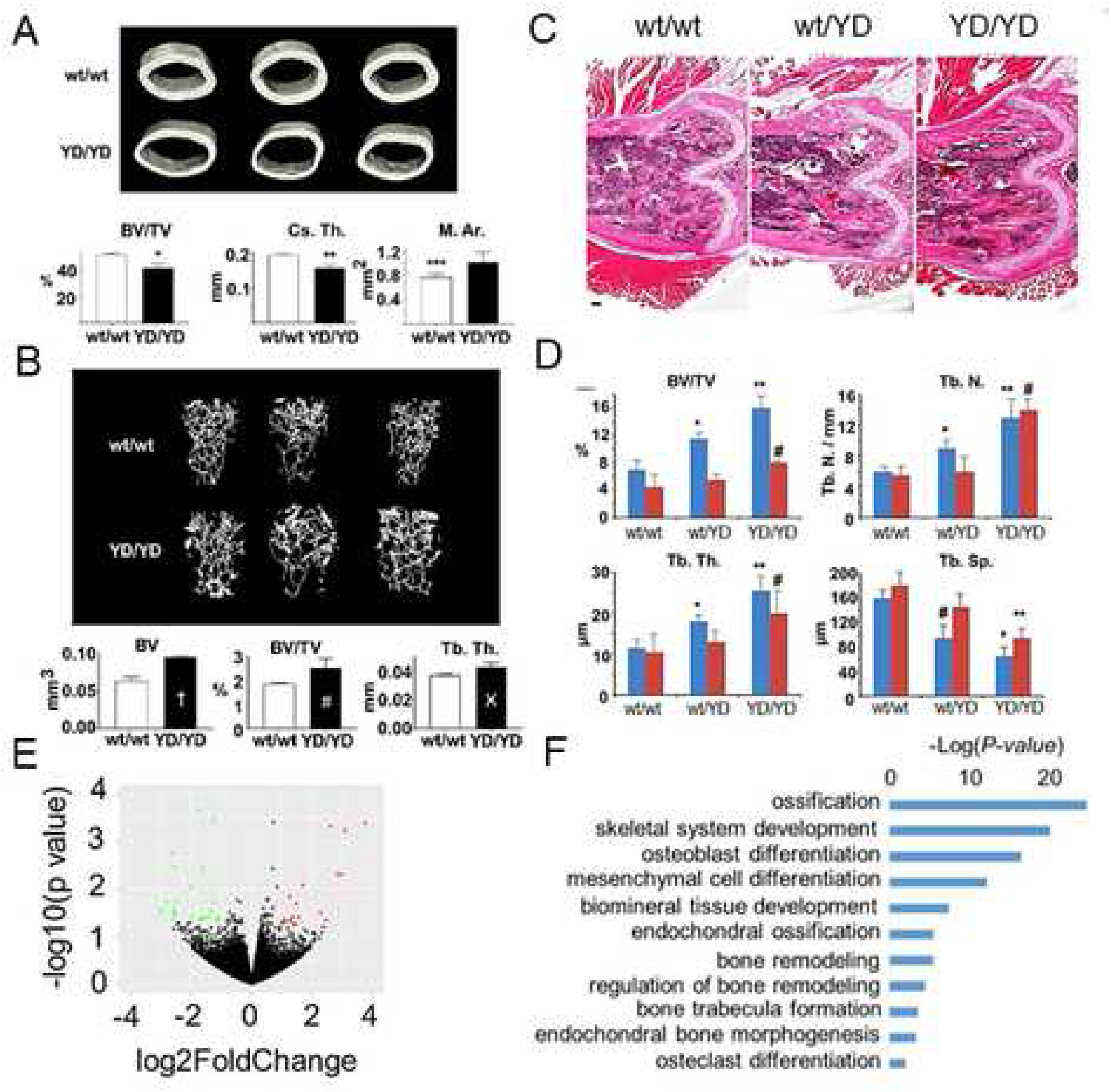
MT1-MMP Y573D mice show decreased cortical bone and increased trabecular bone. **A and B.** MicroCT analysis of cortical (**A**) and trabecular bone (**B**) from *Mmp14^wt/wt^* (wt/wt; n = 3) and *Mmp14^Y573D/Y573D^* (YD/YD; n= 3) 2-month old mice. Upper panels; 3D reconstruction; lower panels: quantitative analysis. **A.** BV/TV: bone volume/tissue volume; *: p = 0.0067; Cs. Th.: cortical thickness; **p = 0.0009; M. Ar.: bone marrow area; *** p = 0.0524. **B.** BV: bone volume;†: p = 0.0115; BV/TV: bone volume/tissue volume; #: p = 0.0366; Tb. Th.: trabecular thickness; Ӿ: p = 0.0290. **C.** Representative sections of the femurs of *Mmp14^wt/wt^* (wt/wt), *Mmp14^Y573D/wt^* (wt/YD) and *Mmp14^Y573D/Y573D^* (YD/YD) 2-month old mice. H&E staining; original magnification: 4X. **D.** Osteomeasure analysis of trabecular bone in femurs (blue bars) and tibiae (red bars) of the mice shown in panel C. Mean ± s.d. of 5 mice/group are shown. BV/TV: bone volume / total volume; *: p = 0.0329; **: p = 0.0012; #: p = 0.0489. Tb. N.: number of trabeculae; *: p = 0.0425; **: p = 0.0021; #: p = 0.0018. Tb. Th.: trabecular thickness; *: p = 0.0478; **: p = 0.0018; #: p = 0.0089. Tb. Sp.: space between trabeculae; *: p = 0.0014; **: p = 0.0019; #: p = 0.0197 (sample vs corresponding wt control). **E.** RNA seq analysis of differential gene expression in bone from *Mmp14^Y573D/Y573D^ vs*. *Mmp14^wt/wt^* mice. Volcano plot representing genes with a significant (p ≤ 0.05) fold change higher than 2 (red dots) or lower than - 2 (green dots). **F.** GO analysis of pathways overrepresented in the bone of *Mmp14^Y573D/Y573D^ vs*. *Mmp14^wt/wt^* mice. See also Figure S2.

Gene expression profiling of bone from *Mmp14^Y573D/Y573D^* mice (Fig. 4 E) showed 76 genes significantly (p ≤ 0.05) up- or downregulated relative to *Mmp14^wt/wt^* littermates. A subset of 17 of these transcripts were upregulated 2-fold or more, and 26 were downregulated 2-fold or more. Consistent with the morphometric analyses, gene ontology analysis showed highly significant enrichment for biological processes related to osteoblast differentiation, bone remodeling, ossification and bone growth (Fig. 4 F).

The knee joints of 2-month old mice showed several abnormalities. Both *Mmp14^Y573D/wt^* and *Mmp14^Y573D/Y573D^* mice displayed marked thinning of articular cartilage (Fig. 5 A top panels, and B) with loss of proteoglycans (Fig. 5 C and D), clustering and cloning of chondrocytes (Fig. 5 A lower panels), classic histologic features of the articular cartilage degeneration associated with human osteoarthritis (OA) and surgical models of OA in mice. The knee cartilage of 2-year old mice also showed fissures, chondrocyte clustering and cloning, abnormalities typical of ageing-associated cartilage degeneration that were not observed in age- and sex-matched *Mmp14^wt/wt^* mice (Fig. 5 E).

**Figure 5.**
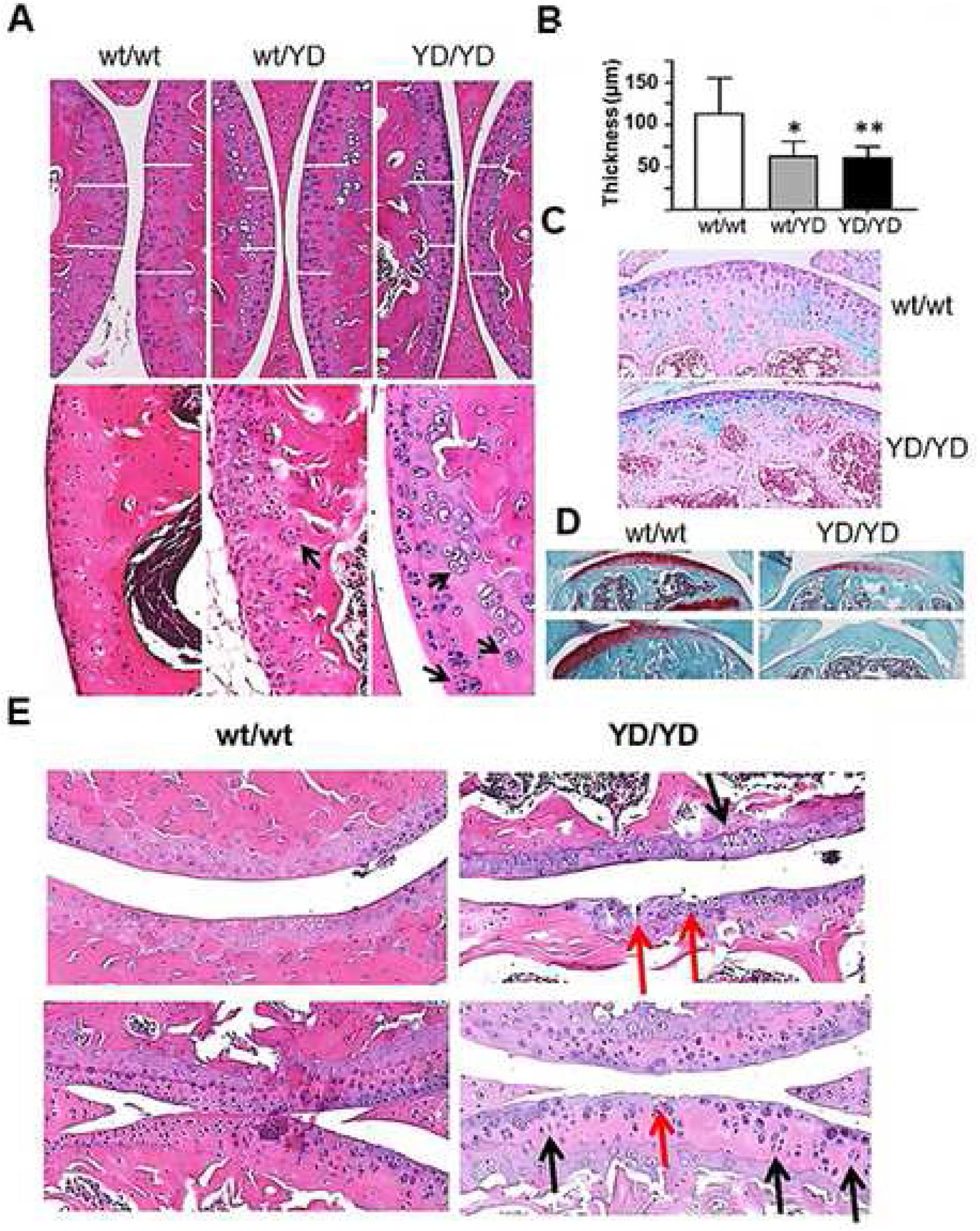
MT1-MMP Y573D mice show reduced thickness and structural abnormalities of articular cartilage. Histological and histochemical analyses of knee joint cartilage from *Mmp14^wt/wt^* (wt/wt), *Mmp14^Y573D/wt^* (wt/YD) and *Mmp14^Y573D/Y573D^* (YD/YD) 2-month old mice. **A.** H&E staining; original magnification: top panels, 10X; bottom panels, 40X. White bars indicate the thickness of the cartilage at comparable locations in the joint. Arrows: chondrocyte cloning. **B.** Thickness of articular cartilage measured by Photoshop as shown in panel A (top panels) on multiple sections of knee joints of 3 mice per genotype. Mean ± s.d. are shown. *: p = 0.0004; **: p = 0.0001. **C.** Alcian blue staining, and **D.** Safranin O staining of sections of knee joints of 2-month old mice with the indicated genotypes. Original magnification: 20X and 10X, respectively. **E.** Histological sections of the shoulder (upper panels) and knee joints (lower panels) of 2-year old *Mmp14^wt/wt^* (wt/wt) and *Mmp14^Y573D/Y573D^* (YD/YD) mice. Red arrows point to cartilage erosion and fissures, and black arrows to chondrocytes oriented orthogonally to the cartilage surface, typical signs of cartilage degeneration. H&E staining; original magnification: 20X.

RNA-Seq analysis of the transcriptome of cartilage from *Mmp14^Y573D/Y573D^* mice (Fig. 6 A) showed significant dysregulation of the expression of 1,549 genes relative to *Mmp14^wt/wt^* mice (p ≤ 0.05). Of these genes, 694 were upregulated 2-fold or more, and 92 were downregulated 2-fold or more. Gene ontology analysis (Fig. 6 B) showed highly significant enrichment for biological processes related to extracellular matrix homeostasis, cartilage development and chondrocyte differentiation. As these biological processes are strongly dysregulated in human OA, we analyzed the transcriptome of *Mmp14^Y573D/Y573D^* cartilage for expression of genes involved in human OA (Fig. 6 C). Gene expression profiling of human OA cartilage has revealed 1,423 genes significantly (p ≤ 0.05) up- or downregulated relative to normal cartilage, 111 of which are strongly up- or downregulated (≥ 2-fold or ≤ 2-fold, respectively) (Geyer et al., 2009; Karlsson et al., 2010). We found that 48 of these 111 OA-associated genes are also strongly regulated in *Mmp14^Y573D/Y573D^* mouse cartilage, including all the genes for collagens and other ECM proteins upregulated in human OA, ECM-degrading proteinases, as well as genes involved in cell metabolism (Figure 6 D). Thus, consistent with its histological features, *Mmp14^Y573D/Y573D^* joint cartilage showed significant gene expression similarity to human OA.

**Figure 6.**
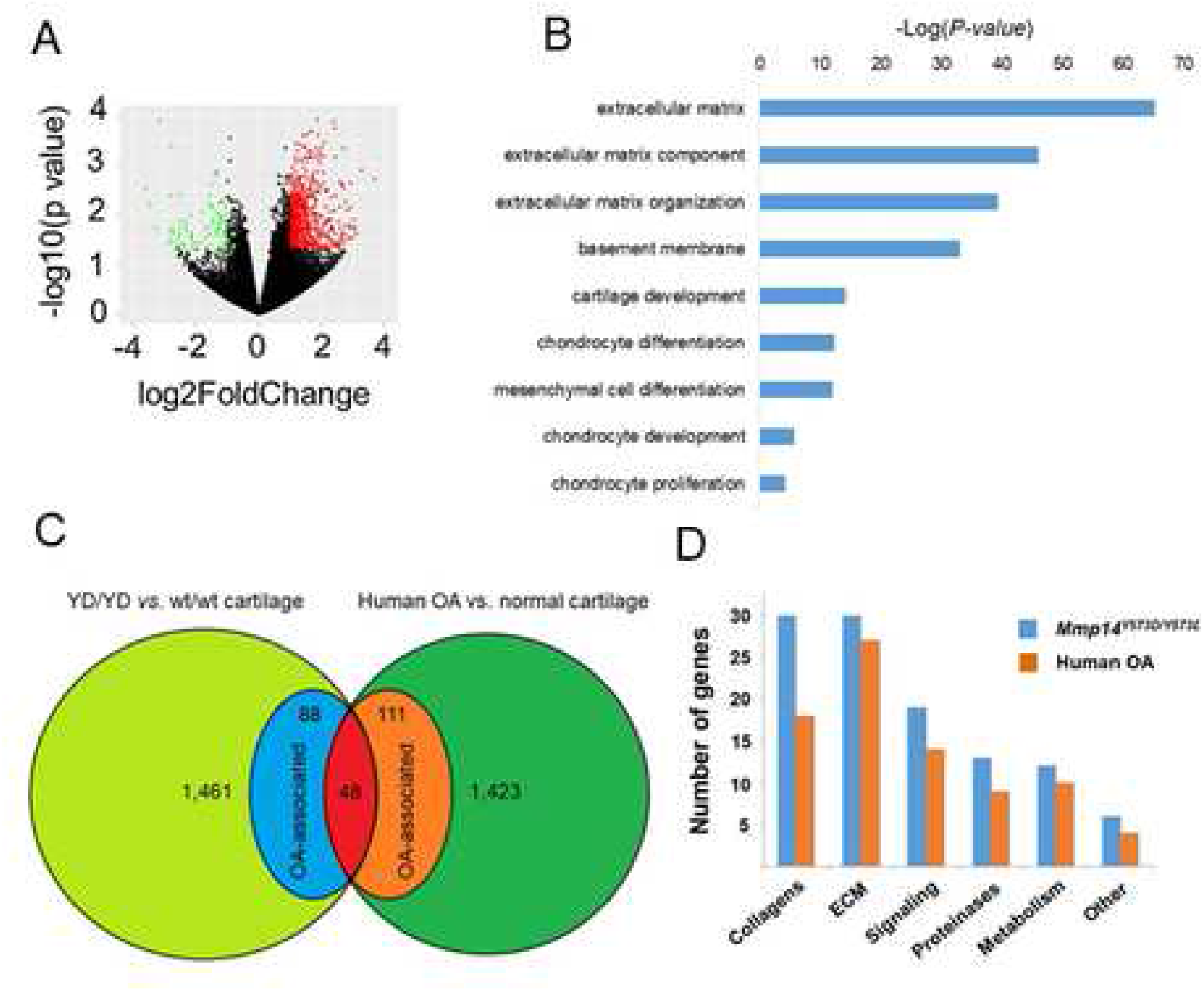
The gene expression profile of the articular cartilage of *Mmp^14Y573D/Y573D^* mice shows similarity to that of human OA. **A.** RNA seq analysis of differential gene expression in articular cartilage from *Mmp14^Y573D/Y573D^ vs*. *Mmp14^wt/wt^* mice. Volcano plot representing genes with a significant (p ≤ 0.05) fold change higher than 2 (red dots) or lower than - 2 (green dots). **B.** GO analysis of pathways overrepresented in the articular cartilage of *Mmp14^Y573D/Y573D^ vs*. *Mmp14^wt/wt^* mice. **C.** Venn diagram of differentially expressed genes in articular cartilage from *Mmp14^Y573D/Y573D^ vs*. *Mmp14^wt/wt^* mice, and human OA cartilage *vs*. normal knee joint cartilage (Geyger et al., 2009; Karlsson et al., 2010). **D.** Gene families upregulated ≥ 2-fold (p ≤ 0.05) in knee joint cartilage from both *Mmp14^Y573D/Y573D^* mice and OA patients.

Sections of the long bones of adult *Mmp14^Y573D/wt^* and *Mmp14^Y573D/Y573D^* mice also showed marked decrease in BM-associated fat relative to *Mmp14^wt/wt^* littermates (Fig. 7 A and B). A similar reduction was apparent in all other fat pads, as evidenced by ~ 50% decrease in body adiposity by DEXA analysis and in the gonadal fat of *Mmp14^Y573D/Y573D^* mice relative to age- and sex-matched *Mmp14^wt/wt^* littermates (Fig. 7 C and D). This effect was observed in mice of both sexes and ages ranging 3 months to 2 years (Figures S3 and S4). Histological analysis of abdominal and subcutaneous WAT from *Mmp14^Y573D/Y573D^* mice revealed marked decrease in the size of adipocytes (Fig. 8 A and C). These findings are consistent with the lipodystrophy of *Mmp14^−/−^* mice (Chun et al., 2006; Chun et al., 2010). In contrast, the brown adipose tissue (BAT) of *Mmp14^Y573D/Y573D^* mice showed pronounced adipocyte hypertrophy, with decreased expression of the characteristic marker of BAT, uncoupling protein-1 (UCP-1; Fig. 8 B and S5).

**Figure 7.**
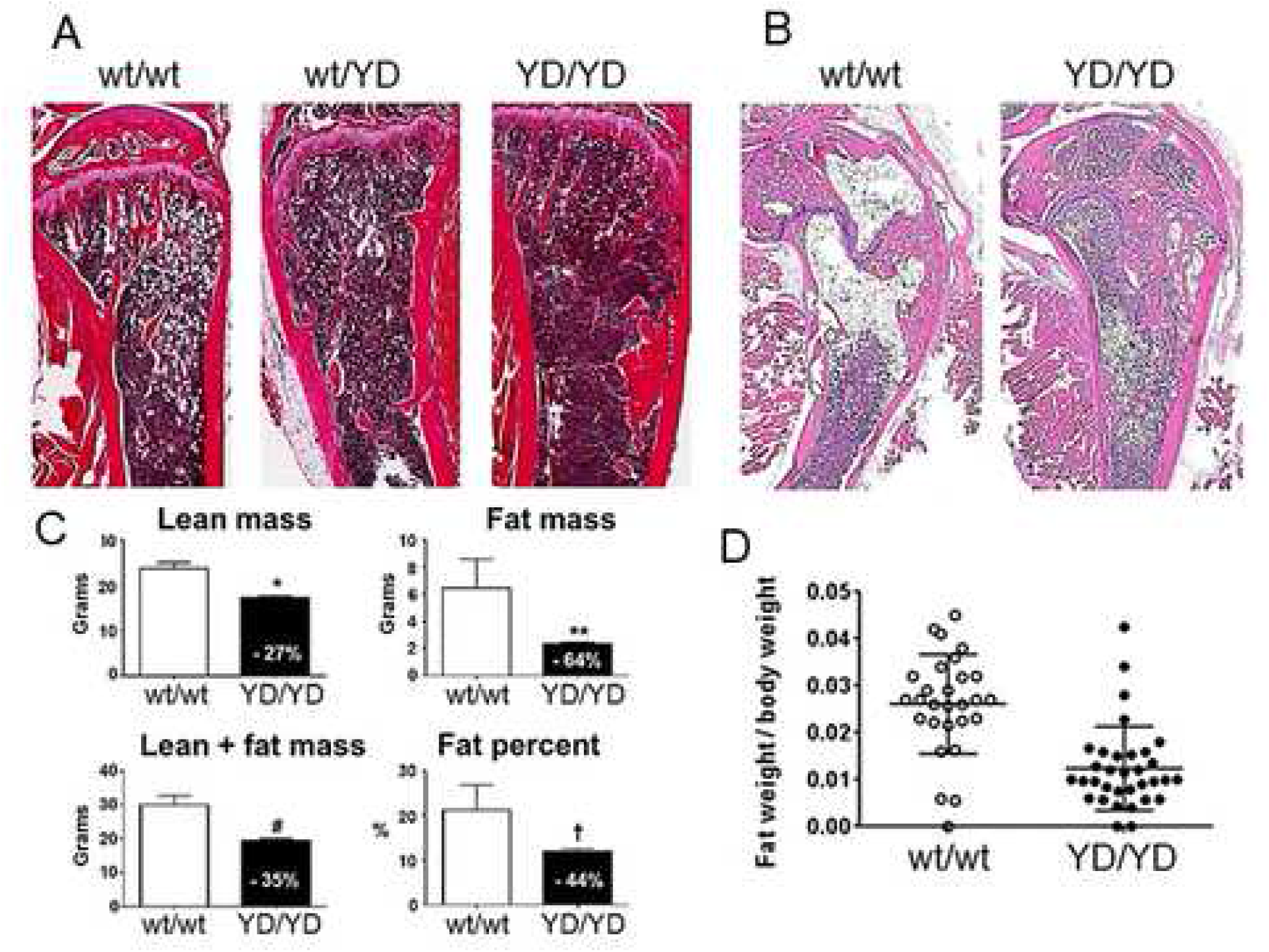
MT1-MMP Y573D mice show decreased white adipose tissue. **A and B.** Sections of the tibias of *Mmp14^wt/wt^* (wt/wt), *Mmp14^wt/Y573D^* (wt/YD) and *Mmp14^Y573D/Y573D^* (YD/YD) 4-month (**A**) or 2-years old mice (**B**). H&E staining; original magnification: 4X. **C.** Dexa scanning analysis of *Mmp14^wt/wt^* (wt/wt; n = 27) and *Mmp14^Y573D/Y573D^* (YD/YD; n = 30) male and female, 7-month old mice. Mean ± s.d. are shown. *: p = 0.0018, **: p = 0.0295, #: p = 0.0177, †: p = 0.0230. Statistically significant percent differences between the two genotypes are indicated in the closed bars. **D**. Weight of abdominal fat normalized to total body weight of 2-, 4- and 7-month old, *Mmp14^wt/wt^* (wt/wt) and *Mmp14^Y573D/Y573D^* (YD/YD) male and female mice; p = 0.0001. See also Figures S3 and S4.

**Figure 8.**
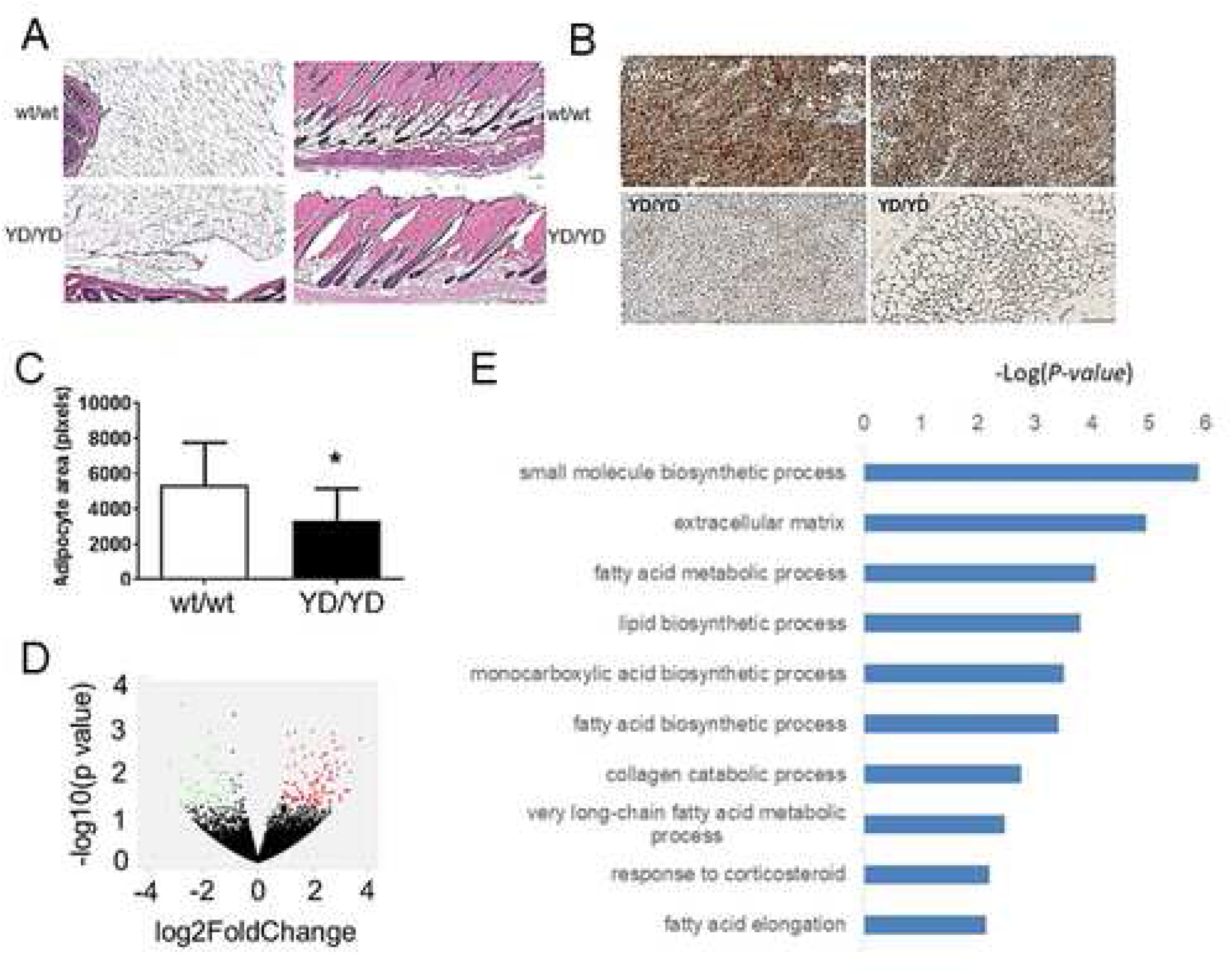
MT1-MMP Y573D mice have hypotrophic white fat and hypertrophic brown fat. **A.** H&E stained sections of periepididymal (left panels; 10X) and subcutaneous fat (right panels; 20X) of *Mmp14^wt/wt^* (wt/wt) and *Mmp14^Y573D/Y573D^* (YD/YD) mice. **B.** Sections of infrascapular brown fat from *Mmp14^wt/wt^* (wt/wt) and *Mmp14^Y573D/Y573D^* (YD/YD) mice, immunostained with antibody to UCP-1. Original magnification: 1.66X, left panels; 20X, right panels. **C.** Adipocyte size of periepididymal fat from *Mmp14^wt/wt^* (wt/wt) and *Mmp14^Y573D/Y573D^* (YD/YD) mice. The area of individual adipocytes was measured by Photoshop using multiple sections (5 mice/genotype) of periepididymal fat similar to those shown in A (left panels). Mean ± s.d. are shown. * p = 0.0003. **D.** RNA seq analysis of differential gene expression white adipose tissue from *Mmp14^Y573D/Y573D^ vs*. *Mmp14^wt/wt^* mice. Volcano plot representing genes with a significant (p ≤ 0.05) fold change higher than 2 (red dots) or lower than - 2 (green dots). **E.** GO analysis of pathways overrepresented in white adipose tissue of *Mmp14^Y573D/Y573D^ vs*. *Mmp14^wt/wt^* mice. See also Figures S3 - S7.

The transcriptome of abdominal WAT from *Mmp14^Y573D/Y573D^* mice (Fig. 8 D) showed significant (p ≤ 0.05) up- or downregulation of the expression of 151 genes, relative to *Mmp14^wt/wt^* mice. Gene ontology analysis (Fig. 8 E) identified highly significant enrichment for biological processes including control of small molecule synthesis, lipid and fatty acid metabolism, response to corticosteroids, and extracellular matrix homeostasis. Some of these pathways are also overrepresented in *Mmp14^+/−^* mice on a high-fat diet (Chun et al., 2010). Moreover, similar to *Mmp14^+/−^* mice, *Mmp14^Y573D/Y573D^* mice were protected from body weight gain induced by high-fat diet (Figure S6 A).

In addition to reduced WAT, *Mmp14^Y573D/Y573D^* mice showed strongly decreased fasting levels of plasma insulin relative to age- and sex-matched *Mmp14^wt/wt^* littermates, with normoglycemia and normal food consumption (Fig. S6 B). *Mmp14^Y573D/Y573D^* and *Mmp14^wt/wt^* mice also had comparable levels of adrenocorticotropic hormone (ACTH), which controls adipose tissue metabolism, and leptin, a hormone secreted predominantly by adipose tissue. Conversely, *Mmp14^Y573D/Y573D^* mice had extremely low serum levels of the inflammatory cytokines interleukin-6 (IL-6), monocyte chemoattractant protein 1/ chemokine (C-C motif) ligand 2 (MCP-1/CCL2), and IL-10 relative to age- and sex-matched *Mmp14^wt/wt^* mice (Fig. S7).

### MT1-MMP Y573D expression alters skeletal stem cell Erk1/2 and Akt signaling, proliferation and apoptosis, and skews differentiation from chondro- and adipogenesis towards osteogenesis

Our previous *in vitro* studies have shown than the Y573D mutation abrogates activation of ERK1/2 and AKT signaling induced by TIMP-2 binding to MT1-MMP (D’Alessio et al., 2008). However, Y573 substitution with D (a negatively charged amino acid) might also mimic phosphotyrosine, and result in constitutive activation of other signaling pathways, including focal adhesion kinase (FAK), whose activation is downregulated in SSC from conditional MT1-MMP knockout mice (Y. Tang et al., 2013). Therefore, we characterized adipose tissue, articular cartilage and bone from *Mmp14^wt/wt^* and *Mmp14^Y573D/Y573D^* mice for activation of Erk1/2, Akt or Fak. The results (Fig. 9 A) showed no significant differences in activation of these signaling pathways between the two genotypes. However, consistent with our previous studies (D’Alessio et al., 2008; Valacca et al., 2015), exogenous TIMP-2 strongly upregulated Erk1/2 and Akt activation in primary fibroblasts from *Mmp14^wt/wt^* mice but had no such effect in *Mmp14^Y573D/Y573D^* cells. Conversely, the levels of active Fak were not increased by exogenous TIMP-2 in either *Mmp14^Y573D/Y573D^* or *Mmp14^wt/wt^* fibroblasts (Fig. 9 B).

**Figure 9.**
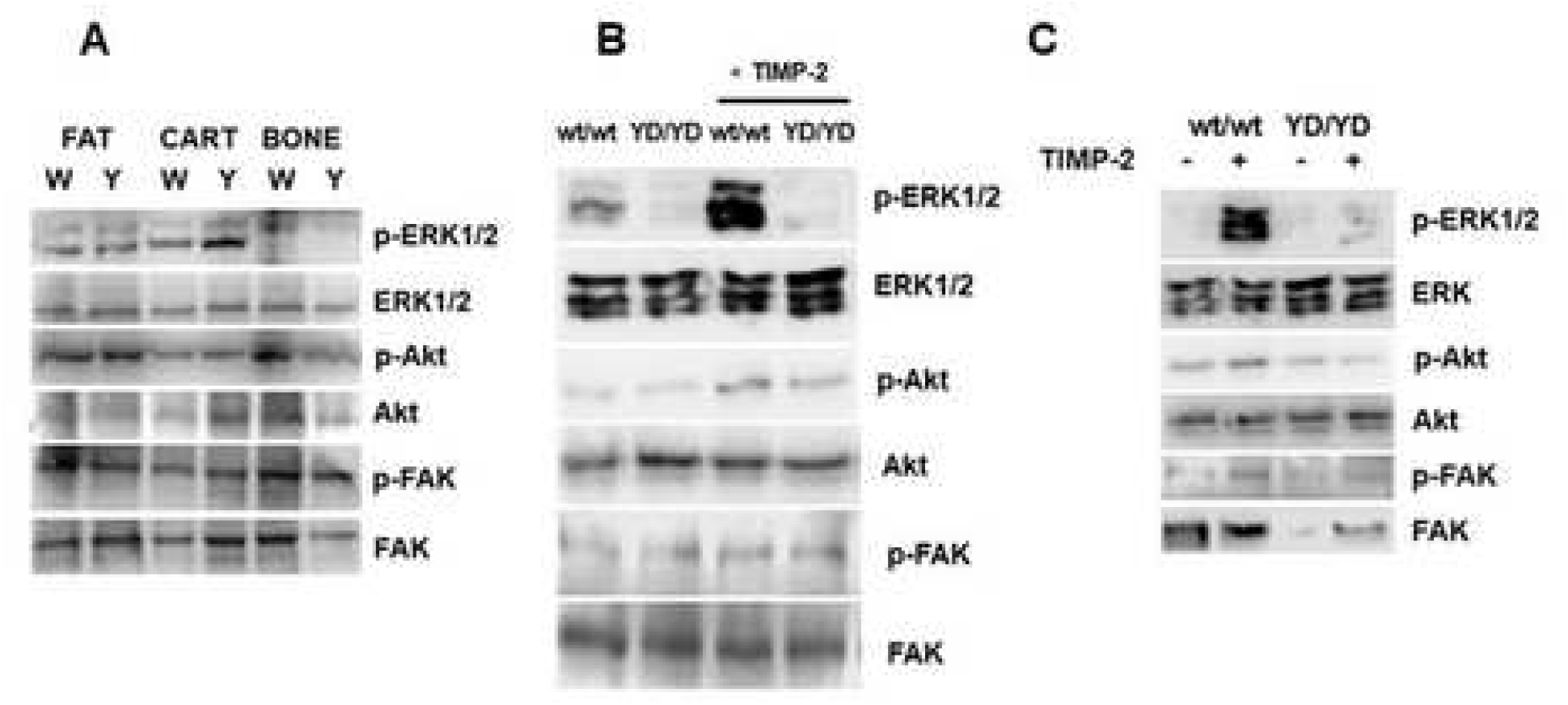
Western blotting analysis of intracellular signaling in tissues and cells from *Mmp14^wt/wt^* and *Mmp14Y573^D/Y573D^* mice. **A.** ERK1/2, Akt and FAK signaling in perigonadal adipose tissue (FAT), knee joint cartilage (CART) and femurs (BONE) of *Mmp14^wt/wt^* (W) and *Mmp14^Y573D/Y573D^* (Y) adult mice. **B and C.** Analysis of ERK1/2, Akt and FAK activation (p-ERK1/2, p-Akt and p-FAK) in primary fibroblasts (B) or BM-derived SSC (C) from *Mmp14^wt/wt^* (wt/wt) and *Mmp14^Y573D/Y573D^* (YD/YD) mice, incubated with human recombinant TIMP-2 (100 ng/ml) or an equivalent volume of control vehicle for 15 min. Total ERK1/2, Akt and FAK protein is shown as a loading control. Equal amounts of cell extract protein were run in two separate gels and transferred onto separate membranes, one of which was probed with antibodies to the phosphorylated forms and the other to the total forms of ERK1/2, Akt and FAK. These experiments were performed two to three times with similar results.

The observation that the phenotype of *Mmp14^Y573D/Y573D^* mice involves abnormalities of bone, cartilage and adipose tissue indicated that the MT1-MMP Y573D mutation might affect the differentiation of SSC, the common progenitor cells of these tissues. Therefore, we isolated SSC from the BM of *Mmp14^wt/wt^* and *Mmp14^Y573D/Y573D^* littermates, and characterized them *in vitro* for Erk1/2, Akt and Fak signaling. The results (Fig. 9 C) showed no significant differences in the basal levels of active Erk1/2, Akt or Fak; however, exogenous TIMP-2 activated Erk1/2 and Akt in *Mmp14^wt/wt^* but not in *Mmp14^Y573D/Y573D^* SSC. Conversely, TIMP-2 did not activate Fak in cells of either genotype.

*Mmp14^Y573D/Y573D^* SSC contained fewer colony-forming units-fibroblasts (CFU-F) than *Mmp14^wt/wt^* SSC (1.5×10^−6^ *vs*. 4.5×10^−6^, respectively), formed much smaller colonies (Fig. 10 A), and showed remarkably lower proliferation and higher apoptosis (Fig. 10 B and C). RNA-Seq analysis (Fig. 10 D) showed 486 genes significantly dysregulated (p ≤ 0.05) in *Mmp14^Y573D/Y573D^ vs*. *Mmp14^wt/wt^* SSC. Gene ontology analysis (Fig. 10 E) revealed highly significant enrichment of genes involved in DNA synthesis, cell cycle regulation and response to DNA damage, indicating dysregulation of cell proliferation and survival, cell functions controlled by Erk1/2 and Akt signaling.

**Figure 10.**
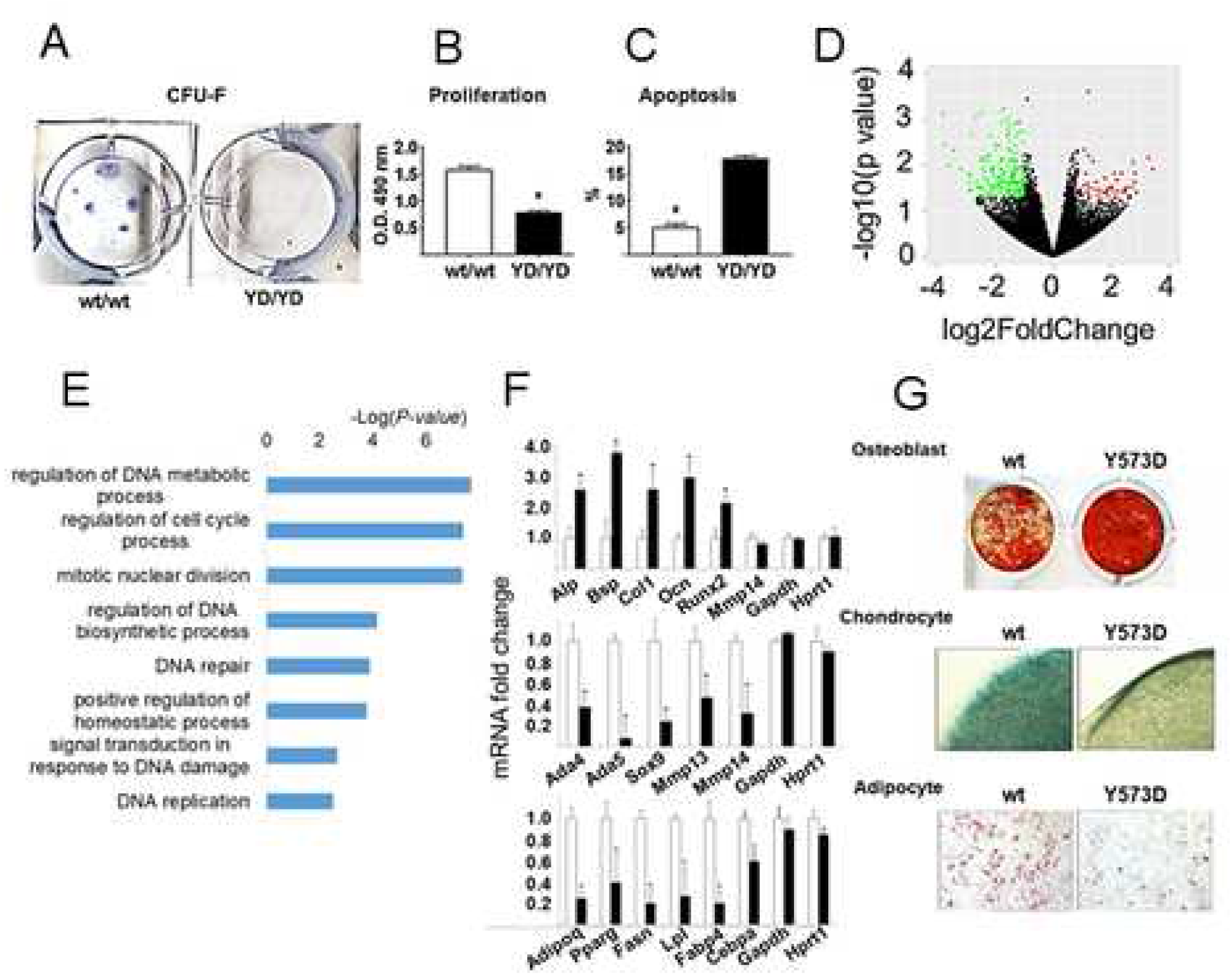
MT1-MMP Y573D expression impairs SSC proliferation and survival, and skews differentiation towards osteogenesis. **A.** CFU-F of *Mmp14^wt/wt^* (wt/wt) and *Mmp14^Y573D/Y573D^* (YD/YD) SSC; Giemsa staining. This experiment was repeated twice with similar results. **B.** MTS assay of cells grown for 48 h. * p = 0.0001. This experiment was repeated three times with similar results. **C.** Percentage of apoptotic nuclei in DAPI stained cells (298 - 320 nuclei). * p = 0.001. This experiment was repeated twice with similar results. **D.** RNA seq analysis of differential gene expression in articular cartilage from *Mmp14^Y573D/Y573D^ vs*. *Mmp14^wt/wt^* mice. Volcano plot representing genes with a significant (p ≤ 0.05) fold change higher than 2 (red dots) or lower than - 2 (green dots). **E.** GO analysis of pathways overrepresented in BM-derived SSC from *Mmp14^wt/wt^* (wt/wt) and *Mmp14^Y573D/Y573D^* (YD/YD) mice. **F.** qPCR analysis of osteoblast (top), chondrocyte (middle) and adipocyte markers (bottom panel) in BM-derived SSC from *Mmp14^wt/wt^* (open bars) and *Mmp14^Y573D/Y573D^* (solid bars) 4-month old littermates, after 17 days in osteoblast or chondrocyte or adipocyte differentiation medium. Shown are mean ± s.d. of data normalized to geomean of GAPDH and HPRT1 for the three cell types. *: p ≤ 0.05. This experiment was repeated twice with similar results. **G.** Differentiation of C3H10T1/2 cells stably transfected with wt or MT1-MMP Y573D cDNA. Top panel, osteogenesis: Alizarin red staining of cells after 21 days in osteoblast differentiation medium. Middle panel, chondrogenesis: Alcian blue staining of cell pellets after 21 days in chondrocyte differentiation medium (original magnification: 4X). Bottom panel, adipogenesis: oil red O staining of cells grown in adipocyte differentiation medium for 7 days (original magnification: 10X). This experiment was repeated three times with similar results.

We then induced BM-derived SSC from *Mmp14^Y573D/Y573D^* and *Mmp14^wt/wt^* littermates to differentiate *in vitro* into the osteoblast, chondrocyte and adipocyte lineages. qPCR analysis of the expression of lineage-specific markers showed a dramatic increase in osteogenesis (Fig. 10 F, top panel), and marked decrease in chondrocyte and adipocyte differentiation (Fig. 10 F, middle and bottom panels, respectively) in *Mmp14^Y573D/Y573D^ vs. Mmp14^wt/wt^* SSC. The expression levels of wt MT1-MMP and MT1-MMP Y573D did not change during osteoblast differentiation, and on day 17 comparable levels of MT1-MMP mRNA were expressed by *Mmp14^wt/wt^* and *Mmp14^Y573D/Y573D^* SSC (Fig. 10 F; top panel). Conversely, in agreement with previous reports (Y. Tang et al., 2013), MT1-MMP expression decreased by ~ 80% during chondrocyte differentiation in both *Mmp14^wt/wt^* and *Mmp14^Y573D/Y573D^* SSC, and on day 17 *Mmp14^Y573D/Y573D^* cells expressed significantly lower MT1-MMP levels than *Mmp14^wt/wt^* SSC (Fig. 10 F, middle panel).

To confirm that the differences in BM-SSC differentiation between *Mmp14^wt/wt^* and *Mmp14^Y573D/Y573D^* mice are mediated by the MT1-MMP Y573D mutation, we transfected wt MT1-MMP or MT1-MMP Y573D cDNA into mouse C3H10T1/2 cells, which are functionally similar to SSC (Q. Q. Tang et al., 2004), and analyzed their capacity to differentiate into osteoblasts, chondrocytes and adipocytes (Fig. 10 G) by lineage-specific staining. Consistent with the results obtained with primary BM-derived SSC, C3H10T1/2 cells transfected with MT1-MMP Y573D showed markedly increased osteogenesis, and decreased chondro- and adipogenesis relative to wt MT1-MMP transfectants (Fig. 10 G). Therefore, in both BM-derived SSC and C3H10T1/2 cells MT1-MMP Y573D expression skewed differentiation towards the osteogenic lineage.

### The bone and fat phenotypes of Mmp14^Y573D/Y573D^ mice are rescued by wt BM transplant

These results indicated that the bone, cartilage and fat phenotypes of *Mmp14^Y573D/Y573D^* mice result from a defect in SSC. Therefore, we hypothesized that these phenotypes could be rescued by transplantation of *Mmp14^wt/wt^* BM; and, *vice versa*, transplantation of *Mmp14^Y573D/Y573D^* BM could transfer the phenotype to *Mmp14^wt/wt^* mice. We therefore transplanted 3- and 5-month old mice with BM from mice of the same age and the opposite genotype or, as controls, from mice of the same age and genotype, and analyzed their phenotypes two months later.

We characterized the trabecular and cortical bone of the femurs of the transplanted mice by microCT analysis (Fig. 11). The results showed that transplantation of *Mmp14^wt/wt^* BM had no effect on the cortical bone of *Mmp14^Y573D/Y573D^* mice transplanted at 3 months or 5 months of age (Fig. 11 A and S8); however, it completely rescued their trabecular bone phenotype (Fig. 11 B and S8). Conversely, transplantation of *Mmp14^Y573D/Y573D^* BM did not transfer the trabecular or cortical bone phenotype to *Mmp14^wt/wt^* mice.

**Figure 11.**
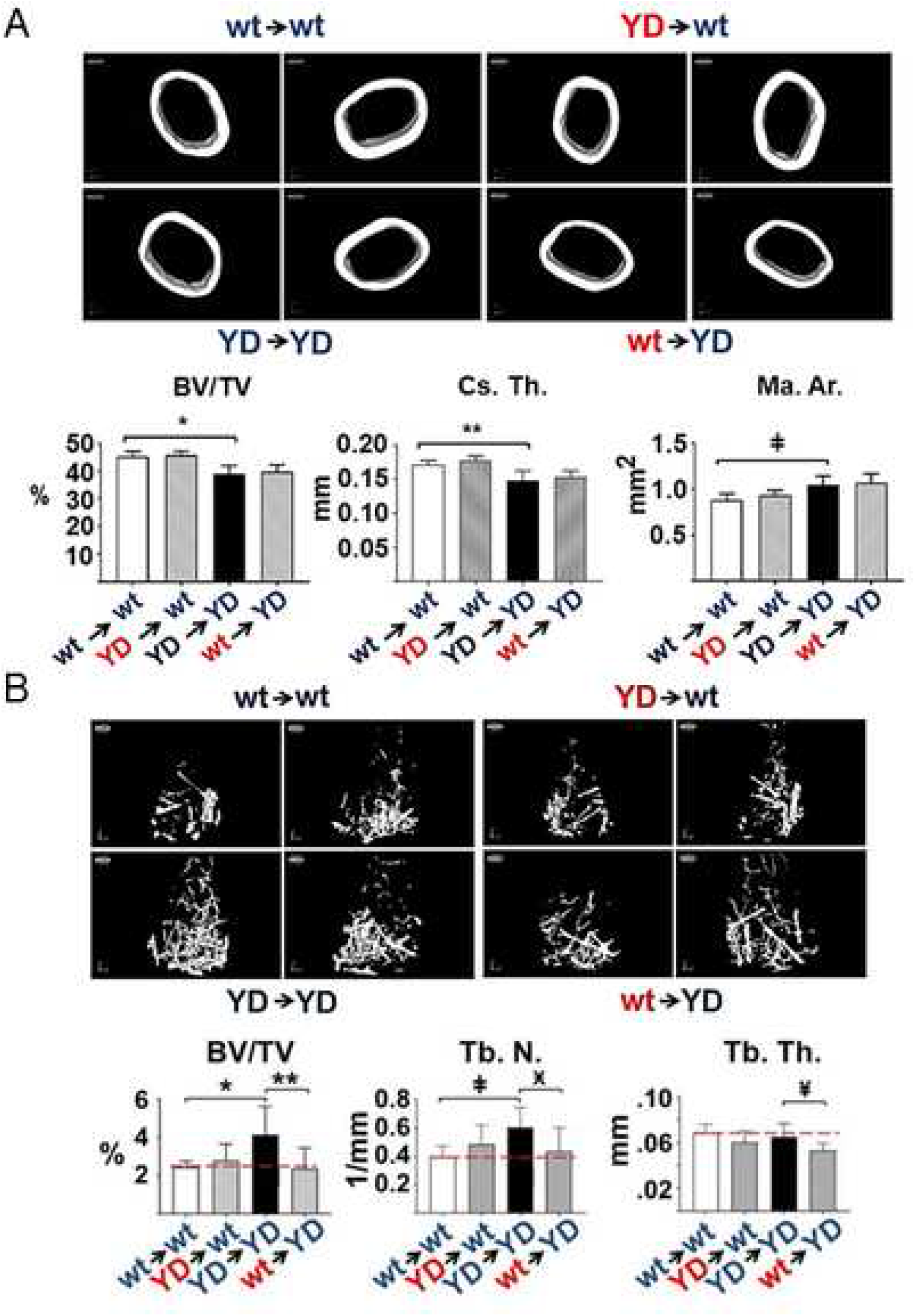
wt BM transplant rescues the trabecular bone phenotype of MT1-MMP Y573D mice. MicroCT 3D reconstruction (upper panels) and quantitative measurements (lower panels) of cortical bone of *Mmp14^wt/wt^* mice transplanted with BM from *Mmp14^Y573D/Y573D^* mice (YD►wt) or *Mmp14^Y573D/Y573^* mice transplanted with *Mmp14^wt/wt^* BM (wt►YD). Mice transplanted with BM of the same genotype (wt►wt and YD►YD) were used as controls. The mice (n = 9-11/group) were transplanted at 3 months of age, and analyzed 2 months later. BV/TV: bone volume normalized to total tissue volume; Cs. Th.: cortical thickness; Ma. Ar.: bone marrow area. The YD►YD samples show ~ 20% reduced cortical bone relative to wt ► wt samples, as expected (Fig. 1 A); wt ► YD or YD► wt BM transplant does not change the phenotype of the recipient. Mean ± s.d. are shown. *: p = 0.0003; **: p = 0.0021; ‡: p = 0.0038. **B.** MicroCT 3D reconstruction (upper panels) and quantitative measurements (lower panels) of trabecular bone of the mice described in A. The YD►YD samples show 35-40% increase in trabecular bone relative to wt ► wt samples, as expected (Fig. 1 B); Mmp14wt/wt BM transplant lowers the amount of trabecular bone in *Mmp14^Y573D/Y573D^* mice to a level similar to that of the wt►wt controls (dotted line). Conversely, BM transplant from *Mmp14^Y573D/Y573D^* mice does not transfer the phenotype to *Mmp14^wt/wt^* mice. BV/TV: bone volume normalized to total tissue volume; Tb. N. trabecular number / mm; Tb. Th.: trabecular thickness. Mean ± s.d. are shown. *: p = 0.047; **: p = 0.0150; ‡: p = 0.0124; Ӿ: p = 0.0397; ¥: p = 0.0117. See also Figures S8 and S9.

We also found that BM transplant induced no significant changes in articular cartilage thickness relative to the original phenotype of the transplant recipients in this short-term study (data not shown).

Similarly to the bone phenotype, *Mmp14^wt/wt^* BM transplant completely rescued the decrease in fat mass of *Mmp14^Y573D/Y573D^* mice. *Mmp14^Y573D/Y573D^* BM did not transfer the adipose tissue phenotype to *Mmp14^wt/wt^* mice (Fig. 12 A). However, consistent with the existence of BM-derived adipocyte precursors able to colonize peripheral WAT and BAT (Crossno et al., 2006), histological analysis of WAT and BAT from transplanted mice (Fig. 12 B and C) showed mixtures of normal and hypotrophic white adipocytes in both *Mmp14^wt/wt^* mice transplanted with *Mmp14^Y573D/Y573D^* BM and *Mmp14^Y573D/Y573D^* mice transplanted with *Mmp14^wt/wt^* BM. Conversely, BM recipients acquired the BAT phenotype of donor mice (Fig. 12 C).

**Figure 12.**
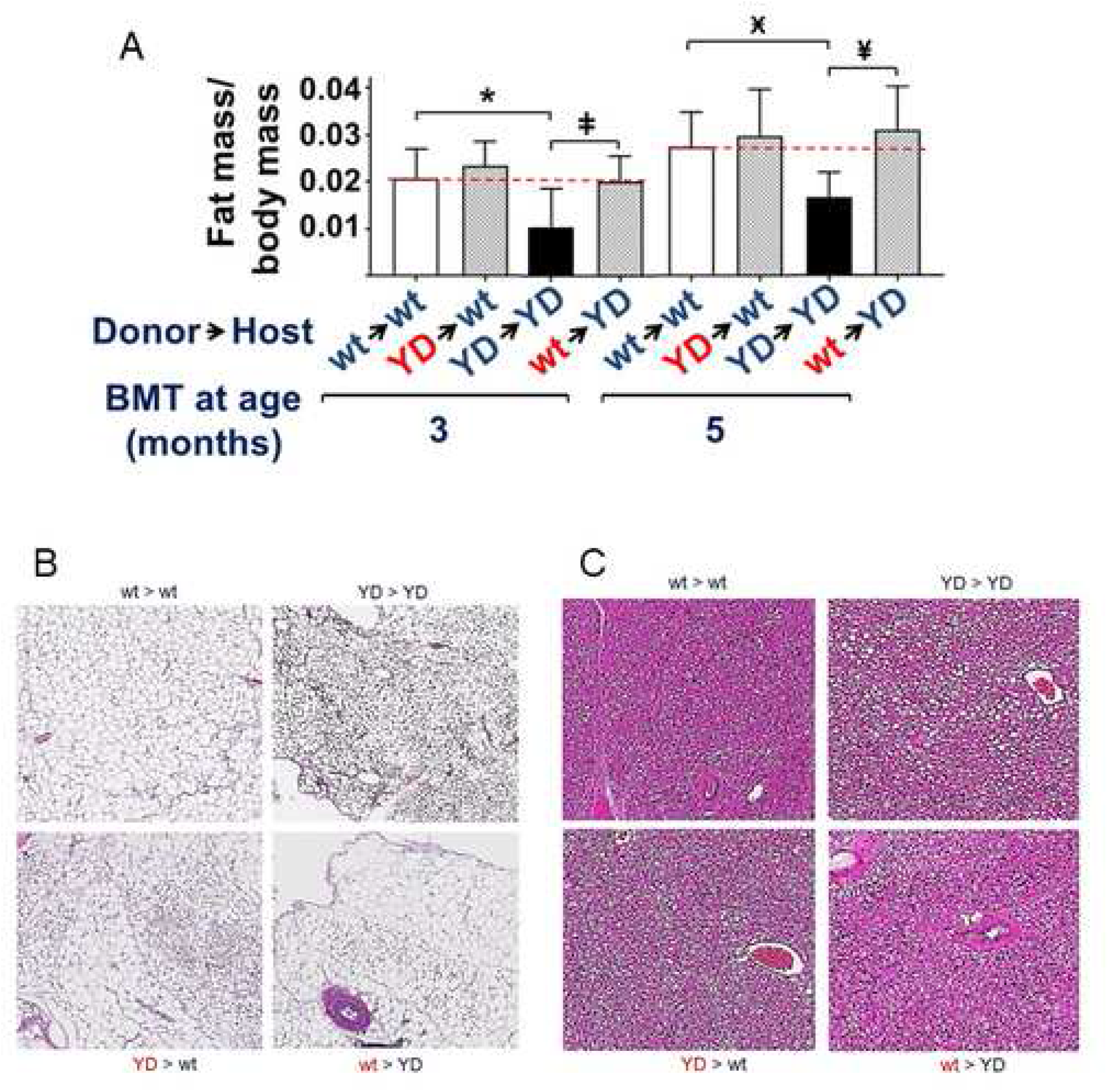
A. wt BM transplant rescues the adipose tissue phenotype of MT1-MMP Y573D mice. Weight of visceral fat normalized to total weight of the mice (n = 9-11/group) described in Fig. 8, transplanted at 3 or 5 months of age and analyzed 2 months later. The YD►YD samples show ~50% less fat than age-matched wt►wt controls, as expected (Fig. 4). *Mmp14^wt/wt^* BM transplant increases the amount of adipose tissue in *Mmp14^Y573D/Y573D^* mice to a level similar to that of the wt►wt controls (dotted line). Conversely, BM transplant from *Mmp14^Y573D/Y573D^* mice does not reduce the fat mass in *Mmp14^wt/wt^* mice. Mean ± s.d. are shown. *: p = 0.006; ‡: p = 0.008; Ӿ: p = 0.003; ¥: p = 0.001. **B and C. WAT and BAT of BM-transplanted mice show a mixture of donor and recipient phenotypes.** Histological sections of periepididymal WAT (**A**) and infrascapular BAT (**B**) of the mice described in Fig. 8. H&E staining; original magnification: A, 10X; B, 20X. The YD►YD samples show hypotrophic white adipocytes (A) and hypertrophic brown adipocytes, as expected (Fig. 5 B and C); wt ► YD or YD► wt BM transplant results in a mixture of normal and hypotrophic white adipocytes (A), and in recipient mice acquiring the BAT phenotype of the donors. See also Figures S8 and S9.

The analysis of BM engraftment, described in Transparent Methods, showed virtually complete replacement of the recipient’s BM with the donors’ BM (Fig. S9). The circulating blood cells of *Mmp14^Y573D/Y573D^* mice displayed no significant abnormalities and numbers comparable to those of *Mmp14^wt/wt^* mice, showing that the MT1-MMP Y573D mutation does not affect hematopoietic cells. Therefore, these results showed that the bone and fat phenotype of transplanted *Mmp14^Y573D/Y573D^* mice resulted from the transfer of SSC, and not from other BM cells.

## DISCUSSION

MT1-MMP plays an important role in the postnatal development of a variety of tissues including bone, cartilage and fat. The striking abnormalities of *Mmp14^−/−^* mice – dwarfism, severe osteopenia, generalized arthritis, fibrosis and lipodystrophy - have been ascribed to impaired collagen turnover, which also plays a fundamental role in SSC differentiation and affects adipocyte development (Chun et al., 2006; Chun & Inoue, 2014; Holmbeck et al., 1999; Zhou et al., 2000). The data presented in this paper show that, in addition to its proteolytic activity, MT1-MMP contributes to bone, cartilage and adipose tissue homeostasis through a proteolysis-independent mechanism mediated by its cytoplasmic domain. This conclusion is based on the following observations.

We have previously shown that MT1-MMP activates ERK1/2 and Akt signaling upon binding of physiological concentrations of TIMP-2 (D’Alessio et al., 2008; Valacca et al., 2015). Signaling is activated by mutant MT1-MMP devoid of proteolytic activity (MT1-MMP E240A) or TIMP-2 lacking MMP inhibitory activity (Ala+ TIMP-2), and is blocked by the Y573D substitution in the MT1-MMP cytoplasmic tail (D’Alessio et al., 2008; Fig. 2 G). Here we showed that the Y573 mutation does not significantly alter the collagenolytic activity of MT1-MMP, or its cleavage of cell membrane proteins *in vivo* and *in vitro* (Figs. 2 - 3). It should also be noted that changes in the levels of membrane proteins cleaved by MT1-MMP result in mouse phenotypes different from those of *Mmp14^Y573D/Y573^* mice. CD44^−/−^ or ADAM9^−/−^ mice have no phenotype (Protin et al., 1999; Weskamp et al., 2002), whereas the genetic deficiency of FGFR2 (cleaved by ADAM9) or Notch1 results in embryonic lethality, and Notch1 haploinsufficiency causes virtually no phenotype (Conlon et al., 1995; Xu et al., 1998). DDR1 deficiency in the mouse results in female infertility and defective lactaction, which we did not observe in *Mmp14^−/−^* mice; conversely, DDR1 haploinsufficiency causes no phenotype (Vogel et al., 2001). Therefore, consistent with our previous findings (D’Alessio et al., 2008; Valacca et al., 2015), our present data show that MT1-MMP activation of Erk1/2 and Akt signaling is independent of its proteolytic activity and mediated by its cytoplasmic domain.

To investigate the physiological significance of our *in vitro* findings, we generated a mouse in which MT1-MMP activation of Erk1/2 and Akt is abrogated by the Y573D mutation. The data presented in this paper show that mice bearing this mutation present phenotypes that partially recapitulate those caused by *Mmp14* deficiency, as well as significantly different phenotypes. The phenotypes of *Mmp14^Y573D/Y573^* mice appear to be milder than those of *Mmp14^−/−^* mice, indicating that both the proteolytic and signaling functions of MT1-MMP are required for normal postnatal development and homeostasis. However, while comparing the phenotypes of *Mmp14^Y573D/Y573^* and *Mmp14^−/−^* mice (Table I) provides information about the relative contribution of the proteolytic and non-proteolytic functions of MT1-MMP to postnatal development and tissue homeostasis, significant limitations must be considered. *Mmp14^−/−^* mice die within 2-3 months of age (Holmbeck & al., 1999), whereas *Mmp14^Y573D/Y573^* mice have a normal life span and their phenotypes become apparent in animals older than 2 months. The global deficiency of *Mmp14* causes severe runting and wasting that ultimately lead to death. It is possible that the dramatic abnormalities of some tissues including cartilage and fat result from indirect effects. For instance, the generalized inflammatory state of *Mmp14^−/−^* mice (Shimizu-Hirota et al., 2012), as opposed to the decreased inflammatory state of *Mmp14^Y573D/Y573D−^* mice, may be the cause of, or contribute to their severe arthritis. Indeed, conditional *Mmp14* knockout in uncommitted SSC (Table I) results in increased thickness of the articular cartilage in 3-month old mice (Y. Tang et al., 2013), in contrast with the reduced thickness and degeneration of *Mmp14^−/−^* and *Mmp14^Y573D/Y573D^* cartilage (Holmbeck & al., 1999).

Furthermore, the relative contribution of MT1-MMP proteolytic and signaling functions to postnatal development and homeostasis may vary at different stages of postnatal life. The lethality of *Mmp14* deficiency limits our understanding of the role of MT1-MMP to relatively early stages of postnatal development; in contrast, the normal life span of *Mmp14^Y573D/Y573^* mice affords studying the role of MT1-MMP in adult animals. Our finding that the phenotype of the *Mmp14^Y573D/Y573^* mouse becomes apparent in adult mice indicates that the proteolytic and the signaling function of MT1-MMP have predominant roles at different stages of postnatal development. The severe phenotype and precocious mortality of the *Mmp14^−/−^* mouse shows that MT1-MMP proteolytic activity has a fundamental role at early stages. Conversely, silencing MT1-MMP signaling is compatible with normal development but results in altered tissue homeostasis in the adult animal.

Our analysis of the articular cartilage of *Mmp14^Y573D/Y573D^* mice showed signs of tissue degeneration similar to, but milder than that of *Mmp14^−/−^* mice. However, whereas *Mmp14^−/−^* mice have severe, acute arthritis (Holmbeck & al., 1999), the articular cartilage of *Mmp14^Y573D/Y573D^* mice shows histological signs and gene expression profile comparable to human OA, a chronic degenerative condition. The striking similarity of the gene expression profile of the articular cartilage *of Mmp14^Y573D/Y573D^* mice to that of human OA raises an interesting point about the role of MT1-MMP in articular cartilage homeostasis and the pathogenesis of OA. MT1-MMP expression decreases during chondrocyte differentiation, and in differentiated chondrocytes is ~ 80% lower than in undifferentiated SSC (Y. Tang et al., 2013). Consistent with this finding, *Mmp14* deficiency in uncommitted SSC results in increased differentiation into chondrocytes and thickening of articular cartilage (Y. Tang et al., 2013), suggesting that MT1-MMP expression contrasts normal chondrocyte development and articular cartilage homeostasis. Indeed, MT1-MMP is upregulated in OA cartilage relative to normal cartilage (Dreier et al., 2004; Kevorkian et al., 2004; Tchetina et al., 2005), indicating that MT1-MMP expression must be strictly controlled for normal cartilage homeostasis. In *Mmp14^Y573D/Y573D^* SSC-derived chondrocytes MT1-MMP expression is lower than in *Mmp14^wt/wt^* mice (Fig. 10 F); however, their cartilage is thinner and presents signs of degeneration, showing that altering MT1-MMP signaling in SSC has a pathological effect even in the presence of low levels of MT1-MMP proteolytic activity. Thus, MT1-MMP signaling is required for articular cartilage homeostasis.

Our phenotypic analysis of *Mmp14^Y573D/Y573D^* mice showed significant abnormalities in WAT, consistent with the phenotype of *Mmp14^−/−^* mice and of mice with conditional *Mmp14* knockout in uncommitted SSC (Y. Tang et al., 2013) (Table 1). However, some differences are noteworthy. In contrast to WAT hypotrophy, we surprisingly found BAT hypertrophy in *Mmp14^Y573D/Y573^* mice, a finding at variance with the normal BAT of *Mmp14^−/−^* mice (Chun et al., 2006). Similarly, BM-associated WAT is decreased in *Mmp14^Y573D/Y573^* mice but increased in mice with conditional knockout of *Mmp14* in uncommitted SSC (Y. Tang et al., 2013) (unfortunately, the severe wasting and early death of *Mmp14^−/−^* mice preclude a reliable assessment of BM-associated fat, which typically develops with aging). However, while the SSC of conditional *Mmp14^−/−^* mice show increased adipocyte differentiation *in vitro* (Y. Tang et al., 2013), the SSC of *Mmp14^Y573D/Y573D^* mice show decreased adipocyte differentiation. MT1-MMP is a fundamental effector of adipocyte growth through collagen degradation, a process required for adipocyte increase in size (Chun et al., 2006; Chun & Inoue, 2014). However, *Mmp14^−/−^* mice have multiple, severe developmental and metabolic defects that can affect adipose tissue development indirectly. Our data show that *in vivo* MT1-MMP contributes to both WAT and BAT homeostasis by a proteolysis-independent mechanism mediated by its cytoplasmic tail. Several studies have shown the involvement of MT1-MMP in adipose tissue homeostasis (Chun et al., 2006; Feinberg et al., 2016; Fenech, Gavrilovic, Malcolm, et al., 2015; Fenech, Gavrilovic, & Turner, 2015), and genetic associations between MT1-MMP and obesity in humans have been reported (Chun et al., 2010). MT1-MMP has also been proposed to control metabolic balance (Mori et al., 2016), a function that could explain the WAT and BAT abnormalities of MT1-MMP Y573D mice. Understanding the proteolysis-independent mechanism of MT1-MMP control of adipose tissue homeostasis can therefore have significant clinical and pharmacological implications.

The cortical bone of *Mmp14^Y573D/Y573^* mice shows a significant decrease in thickness, a phenotype similar to - if milder than - that caused by *Mmp14* deficiency. In contrast, the increased trabecular bone of *Mmp14^Y573D/Y573^* mice contrasts with the severe osteopenia of *Mmp14^−/−^* mice and mice with *Mmp14* knockout in uncommitted SSC (Holmbeck & al., 1999; Y. Tang et al., 2013). This discrepancy between the phenotypic effects of the MT1-MMP Y573D mutation and *Mmp14* deficiency suggests that the proteolytic and signaling functions of MT1-MMP can have opposing roles in bone physiology, and that a balance between the two functions is required for tissue homeostasis. Deletion of the gene abrogates both functions, whereas the MT1-MMP Y573D mutation only affects signaling, altering the balance between ECM proteolysis and intracellular signaling. Moreover, it should be noted again that the bone phenotype of the global or conditional *Mmp14* knockout in SSC can only be observed within the first two-three months of age, whereas the bone abnormalities of *Mmp14^Y573D/Y573^* mice become apparent in animals older than two months. It is possible that the proteolytic and signaling functions of MT1-MMP play different roles in bone modeling (postnatal development) and remodeling (adult life), respectively.

Consistent with their respective phenotypes, the BM-SSC of *Mmp14^Y573D/Y573D^* mice show increased osteoblast differentiation, and decreased chondrocyte and adipocyte differentiation. Our finding that the bone and adipose tissue phenotypes can be rescued by *Mmp14^wt/wt^* BM transplant shows that the *in vivo* effects of the MT1-MMP Y573D mutation result from dysregulation of SSC differentiation. Several considerations can explain the failure of *Mmp14^Y573D/Y573D^* BM to transfer the mutant phenotypes to *Mmp14^wt/wt^* mice, as well as the incapacity of *Mmp14^wt/wt^* BM to rescue the cartilage phenotype of *Mmp14^Y573D/Y573D^* mice. *Mmp14^Y573D/Y573D^* SSC have significantly reduced proliferation and increased apoptosis relative to *Mmp14^wt/wt^* SSC (Fig. 10 A - C). We examined the phenotypes of the transplanted mice 2 months after the transplant. While hematopoietic cells from *Mmp14^Y573D/Y573D^* mice were able to efficiently repopulate the BM of wt mice – indeed, we found no peripheral blood abnormalities in *Mmp14^Y573D/Y573D^* mice – *Mmp14^Y573D/Y573D^* SSC might have required a longer time than wt cells for the phenotypic effects to become apparent. Similarly, BM transplant did not affect the cortical bone phenotype of *Mmp14^Y573D/Y573D^* mice. Cortical bone has a much slower turnover than trabecular bone (Clarke, 2008); therefore a longer time is required for its homeostasis to be altered. The failure of our BM transplant experiments to affect the articular cartilage phenotype is consistent with the absence of vascularization of this tissue. Indeed, no attempts at treating joint cartilage diseases by systemic stem cell administration have thus far been effective (Jevotovsky et al., 2018).

MT1-MMP is constitutively expressed in SSC and its levels are differentially modulated during osteogenic *vs*. chondrogenic/adipogenic differentiation (Y. Tang et al., 2013). The MT1-MMP Y573D mutation and *Mmp14* deficiency have opposing effects on SSC differentiation *in vitro* (Table I). *Mmp14* knockout in uncommitted SSC has no effect on their differentiation in 2D culture; however, it causes decreased osteogenesis and increased chondro- and adipogenesis in 3D collagen gel (Y. Tang et al., 2013). In contrast, in 2D culture MT1-MMP Y573D expression upregulates SSC differentiation into osteoblasts and downregulates chondrocyte and adipocyte differentiation (Fig. 10 F and G). While the *in vitro* differentiation of *Mmp14^Y573D/Y573D^* SSC and SSC with conditional *Mmp14* knockout is consistent with the phenotypes of the respective mice, both mutations fail to fully recapitulate the bone, cartilage and fat abnormalities of the global *Mmp14^−/−^* mouse, which has osteopenia, lipodystrophy and arthritis (Table I). These discrepancies indicate that MT1-MMP controls SSC differentiation by both proteolytic and non-proteolytic mechanisms, and the balance of these two functions is required for normal differentiation. MT1-MMP-mediated ECM degradation modulates mechanosignaling that controls gene expression during SSC differentiation into osteoblasts, and conditional *Mmp14* deficiency in SSC blocks osteogenesis and causes severe osteopenia (Y. Tang et al., 2013). Normal SSC differentiation requires the concerted action of ECM remodeling and intracellular signaling, cell functions modulated by the extracellular environment. *In vitro*, in the presence of abundant ECM - such as in collagen gel culture (Y. Tang et al., 2013) - the proteolytic activity of MT1-MMP is indispensable. Conversely, in the presence of relatively low amounts of ECM – such as in 2D culture – the role of intracellular signaling becomes prevalent. *In vivo,* cell differentiation in the stem cell niche, tissue/organ development and remodeling have different proteolytic and signaling requirements, which are spatially and temporally modulated.

The relative contribution of proteolysis and signaling to MT1-MMP function is also coordinated by extracellular ligands that can inhibit extracellular MT1-MMP proteolytic activity and activate intracellular signaling. This hypothesis is supported by our finding that TIMP-2 binding to MT1-MMP activates ERK1/2 and Akt signaling (D’Alessio et al., 2008; Valacca et al., 2015), as well as by the observation that in the mouse embryo MT1-MMP is temporally and spatially co-expressed with TIMP-2 in the developing skeleton (Apte et al., 1997; Kinoh et al., 1996). *TIMP-2^−/−^* mice do not display the severe phenotype of *Mmp14^−/−^* mice (Z. Wang et al., 2000). We speculate that in these mice signaling can be activated by MT1-MMP binding of TIMP-3 or TIMP-4, as well as a variety of extracellular and transmembrane proteins, including integrins and CD44, that physiologically interact with the MT1-MMP ectodomains (Mori et al., 2002; Zhao et al., 2004).

Thus, the loss of signaling function caused by the Y573D substitution (D’Alessio et al., 2008; Valacca et al., 2015) (Fig. 2 E) can be at the basis of the defects in SSC differentiation and the consequent phenotypes of the *Mmp14^Y573D/Y573D^* mouse. Studies by other groups, as well as our own (unpublished), have shown that, although ERK1/2 signaling is important for osteoblast differentiation (Miraoui et al., 2009; Wu et al., 2015; Xiao et al., 2002), chronic inhibition of ERK1/2 activation results in increased SSC differentiation into osteoblasts (Ge et al., 2007; Higuchi et al., 2002; Nakayama et al., 2003; Schindeler & Little, 2006; Zhang et al., 2012); conversely, inhibition of PI3K/Akt signaling blocks adipo- and chondrogenesis (J. E. Kim & Chen, 2004; Lee et al., 2015; Li & Dong, 2016; Yang et al., 2011). Similarly, the genetic deficiency of Akt (Akt1 and Akt2 double knockout) or ERK1/2 in the mouse results in decreased adiposity and impaired adipogenesis. Conversely, Akt deficiency and conditional knockout of ERK1/2 in osteoprogenitors causes impaired bone development and severe osteopenia (Bost et al., 2005; Cho et al., 2001; J. M. Kim et al., 2019; Peng et al., 2003).

The molecular mechanism that relays signaling from the Y573 residue of the MT1-MMP cytoplasmic tail to the Ras-ERK1/2 and Ras-Akt pathways (D’Alessio et al., 2008; Valacca et al., 2015), remains to be investigated. The adaptor protein p130Cas, which binds to the cytoplasmic tail of MT1-MMP by a mechanism involving Y573 (Gonzalo et al., 2010; Y. Wang & McNiven, 2012), recruits Src and/or focal adhesion kinase (FAK), which activate Ras (Bunda et al., 2014; Schlaepfer & Hunter, 1997). We speculate that TIMP-2 binding to MT1-MMP triggers the assembly of the p130Cas/Src/FAK complex at the MT1-MMP tail, and that this effect is abrogated by the Y573D substitution, which thus prevents the downstream activation of ERK1/2 and AKT signaling. In addition, Y573 could control intracellular signaling by modulating cytoplasmic tail interactions with a variety of transmembrane or membrane-bound proteins including caveolin-1 (Annabi et al., 2001; Galvez et al., 2002; Galvez et al., 2004; Labrecque et al., 2004) and ß1 integrins or growth factor receptors (Langlois et al., 2007). The cytoplasmic tail of MT1-MMP also controls energy production by activating hypoxia-inducible factor-1 (HIF-1) by a non-proteolytic mechanism (Sakamoto & Seiki, 2010).

In conclusion, our findings provide the first *in vivo* evidence for an important role of MT1-MMP mediated, proteolysis-independent signaling in postnatal development and tissue homeostasis. Understanding the relative contribution of the proteolytic and signaling functions of MT1-MMP to the control of metabolic processes that affect a variety of tissues and organs will require the development of additional genetically engineered mouse models, as well as the molecular dissection of extracellular and intracellular components of the MT1-MMP signaling mechanism. The knowledge obtained from these studies can increase our understanding of the pathogenesis of important diseases that affect bone, cartilage and adipose tissue homeostasis, such as osteopenia, osteoarthritis and obesity, and potentially direct the design of pharmacological tools for their treatment.

## LIMITATIONS OF THE STUDY

Our results show that the Y573D mutation in the MT1-MMP cytoplasmic tail abrogates TIMP-2 induced activation of Erk1/2 and Akt in SSC, suggesting that this effect is responsible for the defects in SSC differentiation and the development of the MT1-MMP Y573D mouse phenotypes. However, this mutation may also block or activate other signaling pathways. Future studies are warranted to understand the molecular mechanism by which the Y573 residue of the MT1-MMP cytoplasmic tail activates intracellular signaling that controls tissue homeostasis.

## Resource Availability

### Lead Contact

Paolo Mignatti, NYU School of Medicine, Department of Medicine, Division of Rheumatology, 301 East 17th Street, Suite 1612A, New York, NY 10003, USA.

### Materials Availability

The Mmp14^Y573D/Y573D^ mouse is available upon request to the lead contact.

### Data and Code Availability

The RNA-seq raw data are available at doi:10.5061/dryad.s4mw6m950.

## ACKNOWLEDGMENTS

This work was supported by NIH grants R01 CA136715, R01CA136715-05S1 and R21 AG033735 (to P.M.) and in part by R21 AR069240-01A1 (to S.A.). We gratefully acknowledge the precious collaboration of the Rodent Genetic Engineering, High-Throughput Biology Laboratory, Experimental Pathology Research Laboratory, and Genome Technology Center of NYU School of Medicine, which are partially supported by NIH grant P30CA016087 to the Laura and Isaac Perlmutter Cancer Center and the Shared Instrumentation Grant S10 OD021747, and the MicroCT Core of NYU College of Dentistry, supported by NIH grant S10 OD010751 to Dr. Nicola C. Partridge.

## AUTHOR CONTRIBUTIONS

Conceptualization, P.M.; Methodology, P.M., M.A., X.Z., B.B.D., S.Y. and S.A.; Validation, P.M., M.A.; Formal Analysis: P.M., M.A. and C.L.; Investigation, M.A., C.L., X.Z., T.H, C.A., C.V., S.Z., S.M., E.V., Q.Y., V.K. and S.Y.; Data Curation, P.M.; Writing – Original Draft, P.M.; Writing – Review & Editing, P.M., M.A., S.Y. and S.A.; Visualization, P.M., M.A. and S.A.; Supervision, P.M. and M.A.; Project Administration, P.M.; Funding Acquisition, P.M. and S.A.

## DECLARATION OF INTERESTS

The authors declare no competing interests.

---

**Figure S1.**
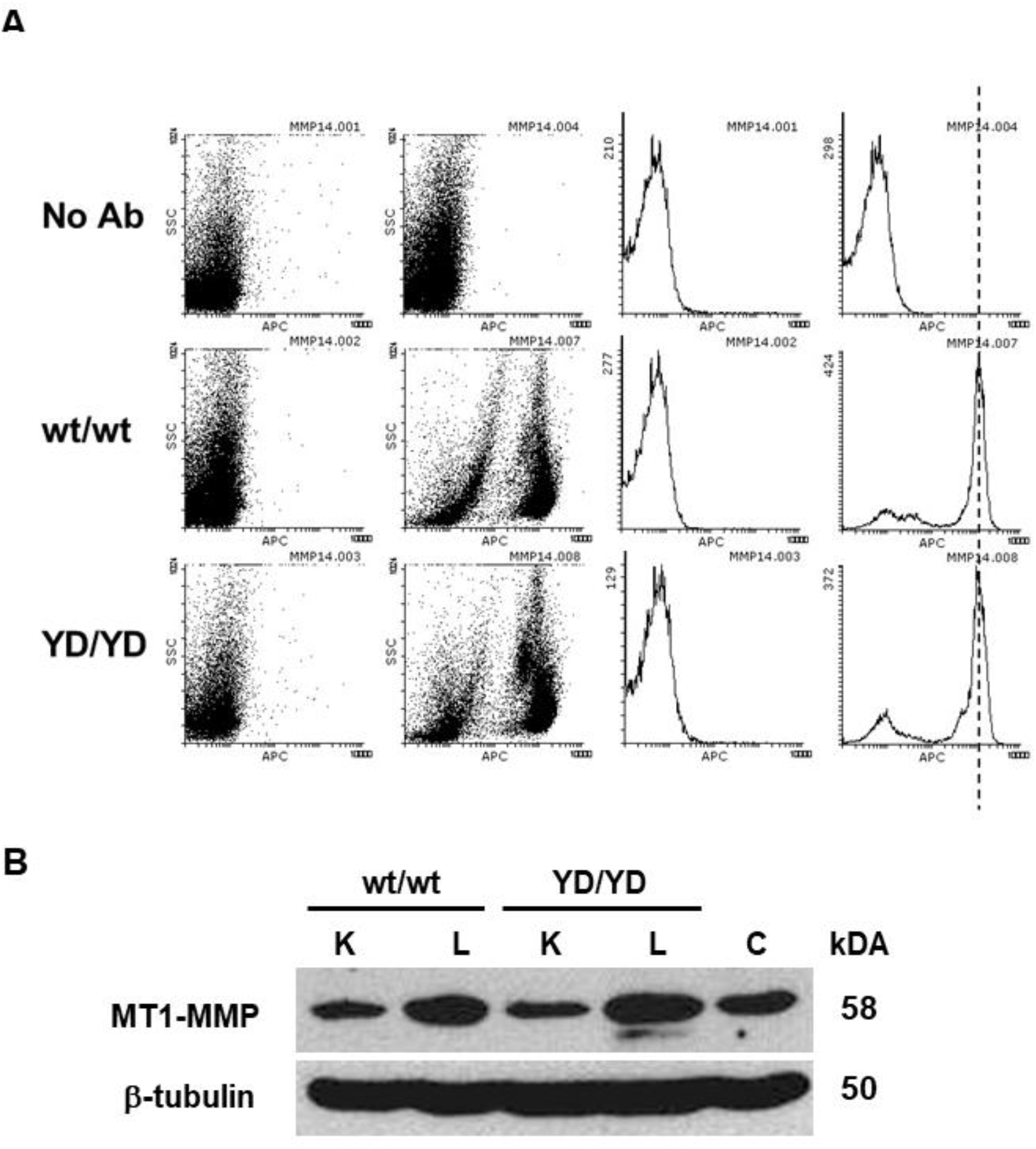
Analysis of wt MT1-MMP and MT1-MMP Y573D expression *in vivo*. Related to Figures 2 and 3. **A**. Flow cytometric analysis of MT1-MMP on the surface of peripheral blood monocytes from *Mmp14*^*wt/wt*^ (wt/wt) and *Mmp14*^*Y573D/Y573D*^ (YD/YD) male mice. **B.** Western blotting analysis of MT1-MMP in protein extracts of lungs and kidneys from 3-month old *Mmp14*^*wt/wt*^ (wt/wt) and *Mmp14*^*Y573D/Y573D*^ (YD/YD) male mice. Tissues from three littermates per genotype were pooled for protein extraction.

**Figure S2.**
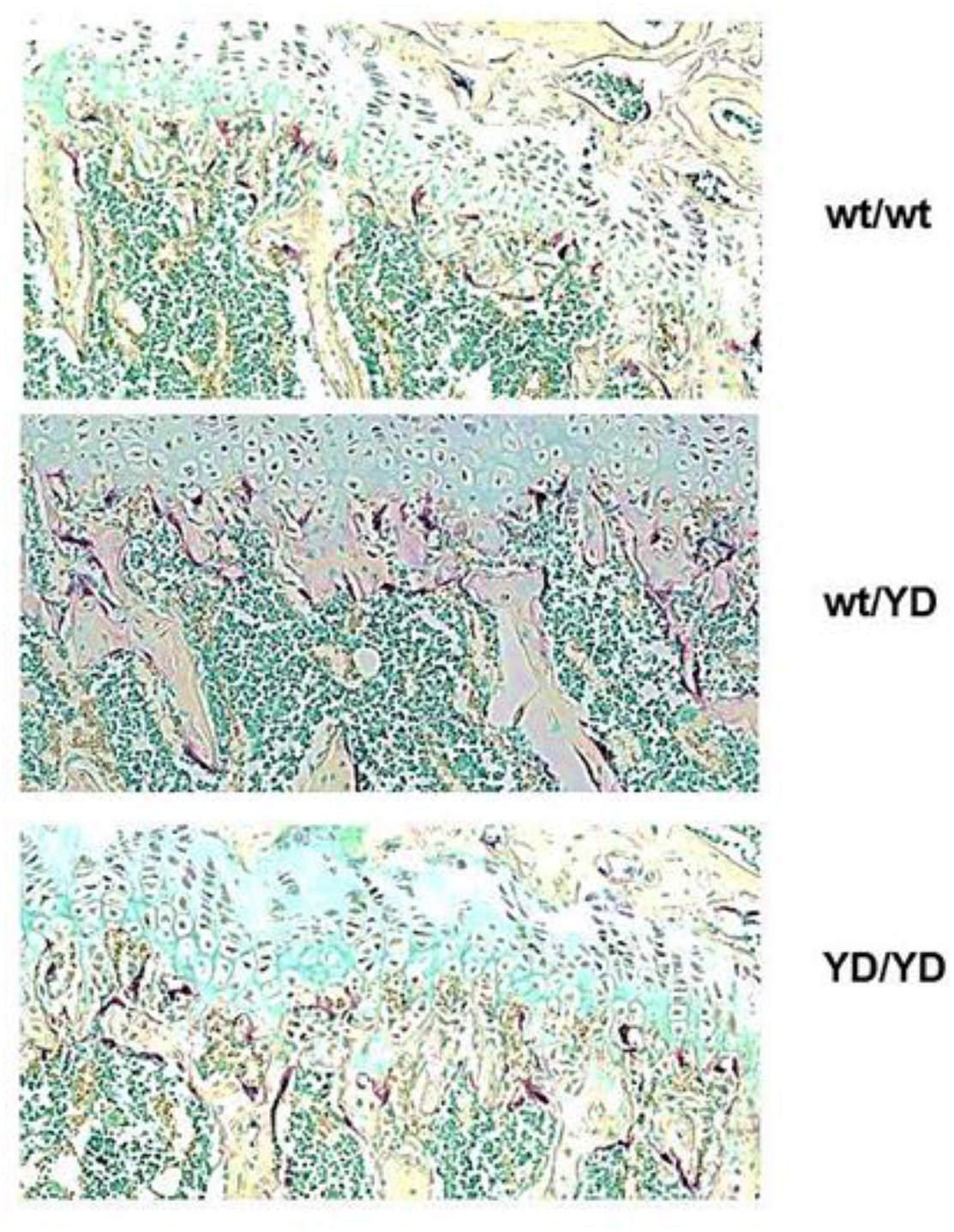
Analysis of osteoclasts in the trabecular bone of *Mmp14^wt/wt^*, *Mmp14^wt/Y573D^* and *Mmp14^Y573D/Y573D^* mice. Related to figure 4. Tartrate-resistant acid phosphatase (TRAP) staining of osteoclasts (dark red cells) in sections of the femurs of *Mmp14*^*wt/wt*^ (wt/wt), *Mmp14*^*wt/Y573D*^ (wt/YD) and *Mmp14*^*Y573D/Y573D*^ (YD/YD) 2-month old mice. Original magnification: 4X

**Figure S3.**
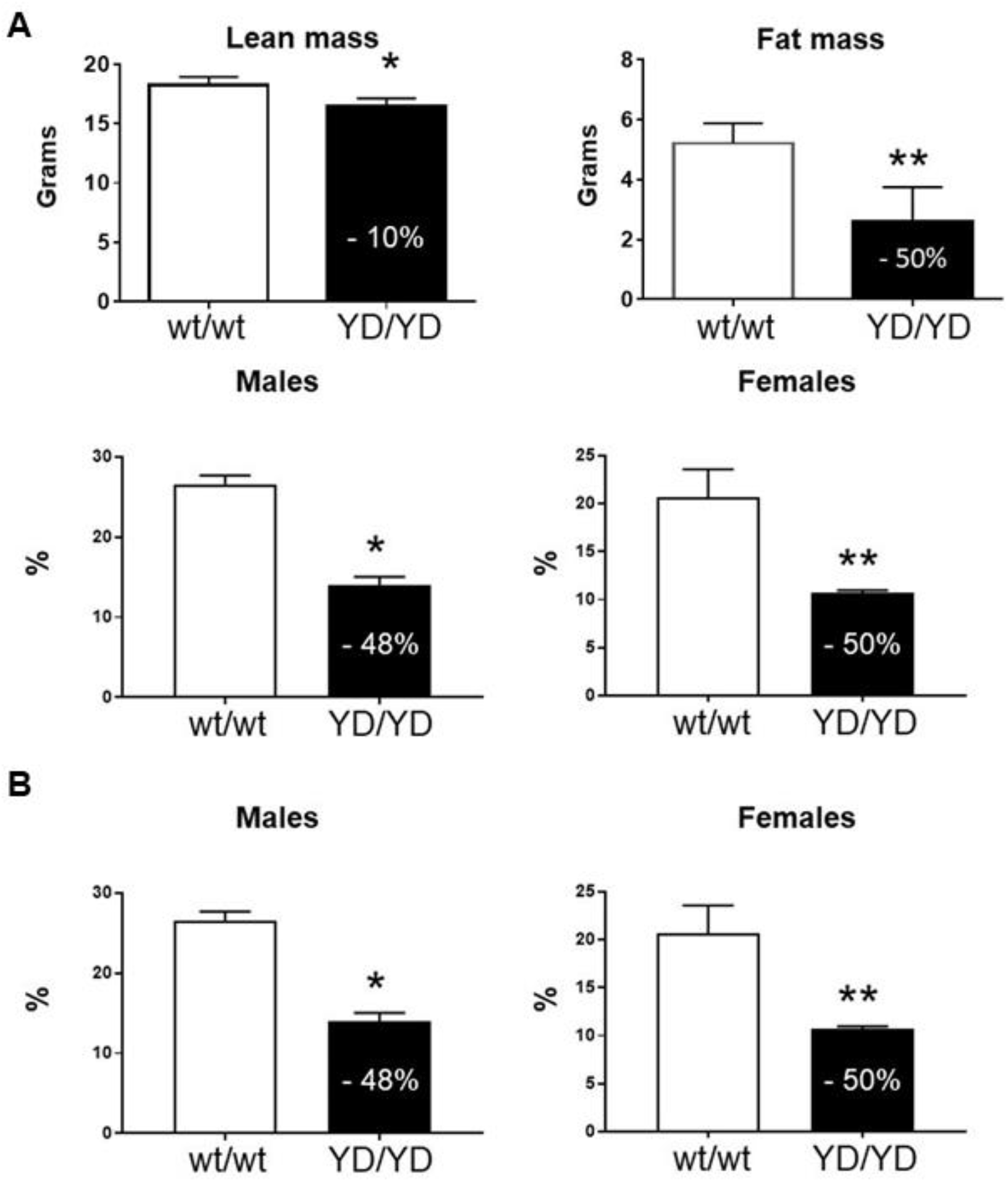
Analysis of adipose tissue in *Mmp14^wt/wt^* and *Mmp14^Y573D/Y573D^* mice. Related to Figures 7 and 8. **A.** DEXA scanning analysis of lean and fat body mass of 4-month old *Mmp14*^*wt/wt*^ (wt/wt; n = 10) and *Mmp14*^*Y573D/Y573D*^ (YD/YD; n = 10) female mice. Mean ± s.d. are shown. Statistically significant percent differences between the two genotypes are indicated in the closed bars. *: p = 0.0121; **: p = 0.0060; #: p = 0.0093; ¶: p = 0.0124. **B.** DEXA scanning analysis of adipose tissue in 10-weeks old *Mmp14*^*wt/wt*^ (wt/wt; n = 10) and *Mmp14*^*Y573D/Y573D*^ (YD/YD; n = 10) male or female mice. Mean ± s.d. are shown. Statistically significant percent differences between the two genotypes are indicated in the closed bars. *: p = 0.0001; **: p = 0.0044.

**Figure S4.**
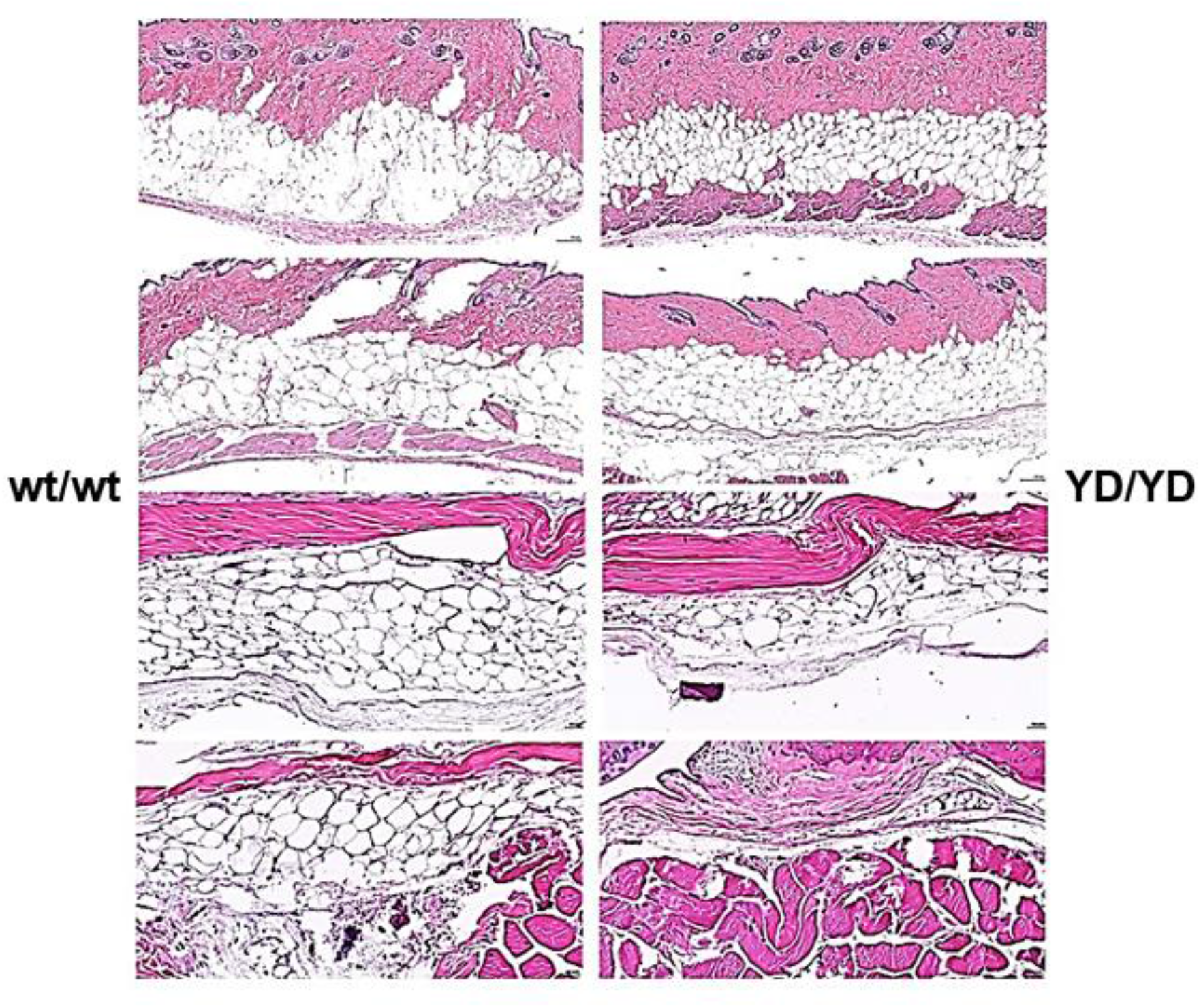
Analysis of subcutaneous fat in *Mmp14^wt/wt^* and *Mmp14^Y573D/Y573D^* mice. Related to Figures 7 and 8. Histological sections of subcutaneous fat in *Mmp14*^*wt/wt*^ (wt/wt; left column) and *Mmp14*^*Y573D/Y573D*^ mice (YD/YD; right column). H&E staining; original magnification: 20X. Age- and sex-matched mice are shown on the same row. *Mmp14*^*Y573D/Y573D*^ mice show hypotrophic adipocytes and reduced thickness of the adipose layer.

**Figure S5.**
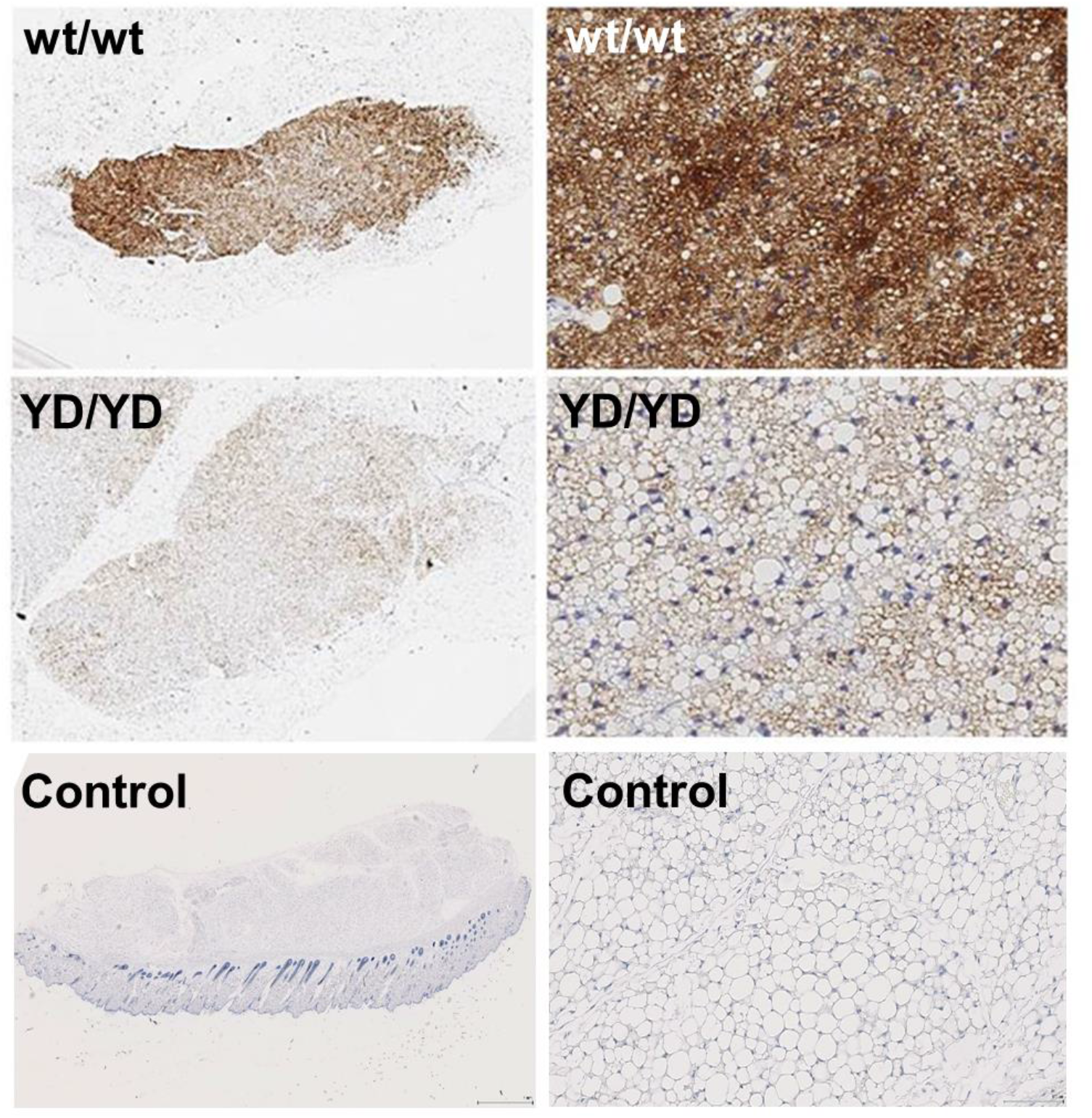
Analysis of brown fat in *Mmp14^wt/wt^* and *Mmp14^Y573D/Y573D^* mice. Related to Figures 7 and 8. Histological sections of brown fat from *Mmp14*^*wt/wt*^ (wt/wt) and *Mmp14*^*Y573D/Y573D*^ mice (YD/YD), immunostained with antibody to UCP-1. Control: section of subcutaneous white fat from a wt mouse, immunostained with UCP-1 antibody using the same protocol as for the brown fat sections. Original magnification: 1.66X, left panels; 20X, right panels.

**Figure S6.**
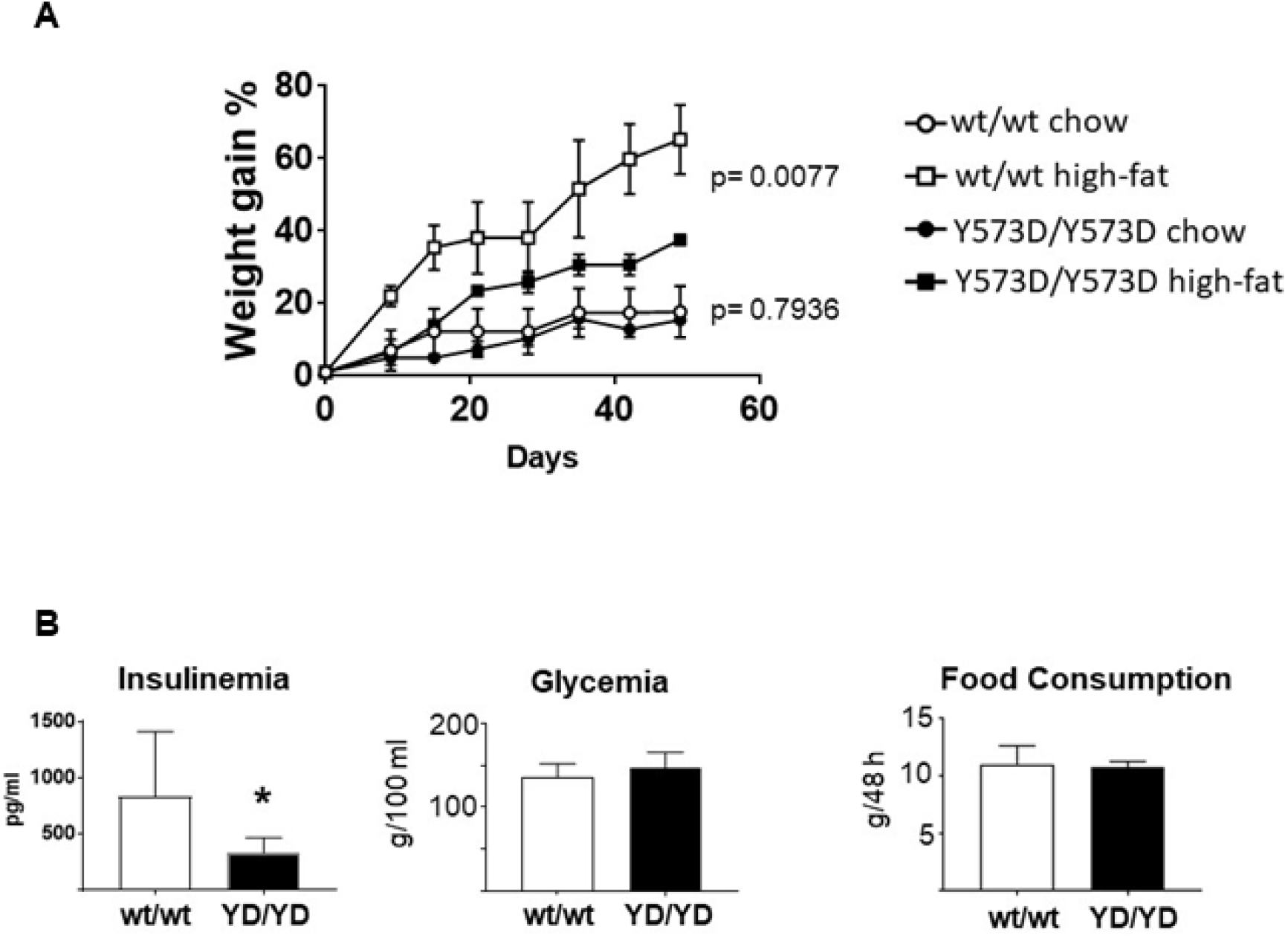
Metabolic analysis of *Mmp14^wt/wt^* and *Mmp14^Y573D/Y573D^* mice. Related to Figures 7 and 8. **A.** Weight gain analysis of 6-week old *Mmp14*^*wt/wt*^ (wt/wt; n = 2) and *Mmp14*^*Y573D/Y573D*^ (YD/YD; n = 2) littermates fed normal chow or a high-fat diet. The regression curves and the statistical significance of their differences were calculated with GraphPad Prism, Version 7.05. **B.** Fasting serum levels of insulin and glucose in, and food consumption by, *Mmp14*^*wt/wt*^ (wt/wt; n = 8) and *Mmp14*^*Y573D/Y573D*^ (YD/YD; n = 8) mice. The graph of insulinemia shows the same data as in Fig. S7, and is shown here for comparison with glycemia. *: p = 0.0203.

**Figure S7.**
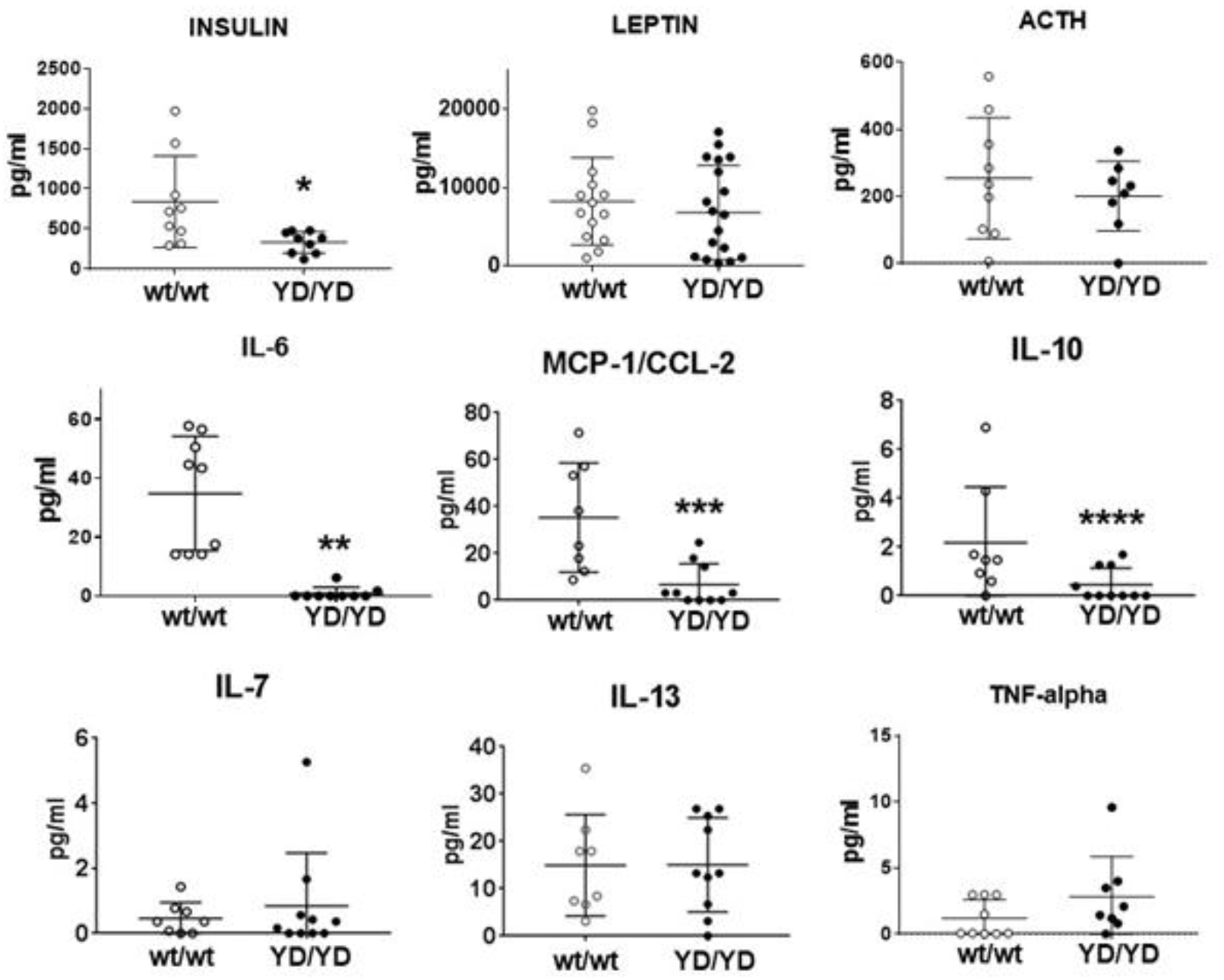
*Mmp14^Y573D/Y573D^* mice have low levels of insulin and inflammatory cytokines. Related to Figures 7 and 8. Analysis of circulating hormones and cytokines in 4-month old *Mmp14*^*wt/wt*^ (wt/wt) and *Mmp14*^*Y573D/Y573D*^ (YD/YD) mice. Mean ± s.d. are shown. *: p = 0.023; **: p = 0.0001; ***: p = 0.0024; ****: p = 0.0384.

**Figure S8.**
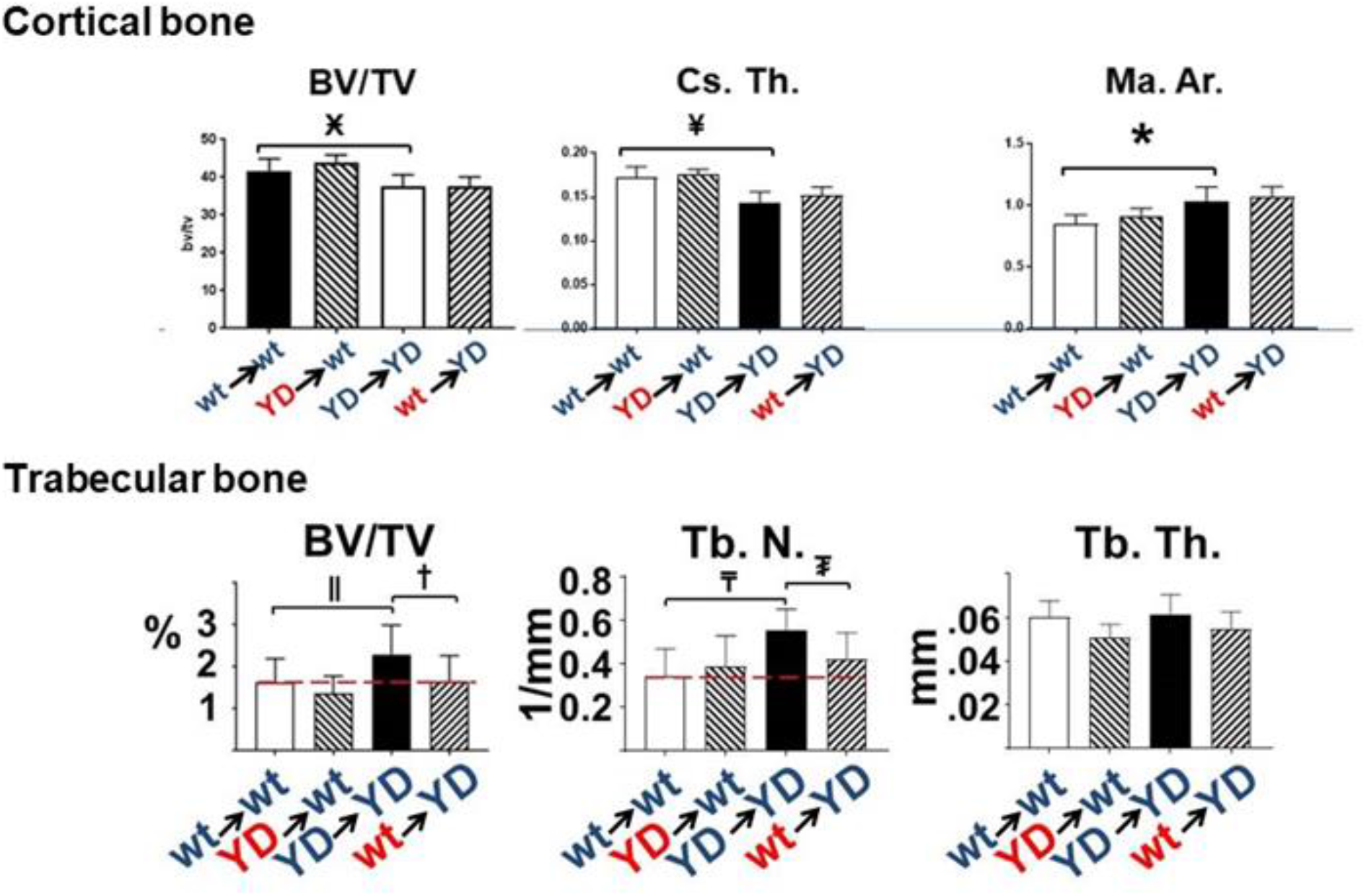
BM transplant rescues the trabecular bone phenotype of MT1-MMP Y573D mice. Related Figure 11. Quantitative microCT analysis of cortical and trabecular bone of 5-month old mice (n = 9-11/group) transplanted as described in the legend to Fig. 11. The mice (n = 9-11/group) were transplanted at 5 months of age, and analyzed 2 months later. The results are comparable to those presented in Figs. 13 and 14. Ӿ: p = 0.002; ¥: p = 0.0002; *: p = 0.005; ǁ: p = 0.0323; †: p = 0.0362; ₸: p = 0.0025; ₮: p = 0.0162.

**Figure S9.**
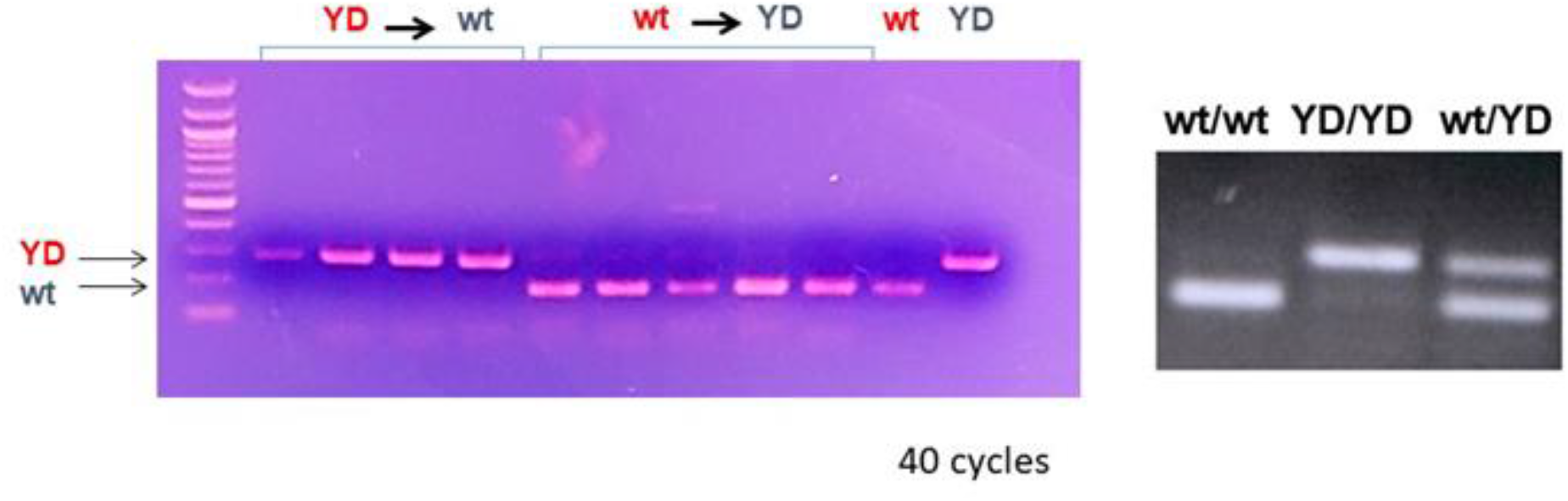
Transplanted bone marrow from *Mmp14^wt/wt^* mice replaces the bone marrow of *Mmp14^Y573D/Y573D^* mice. Related to Figures 11 and 12. PCR genotyping of BM from mice transplanted with BM as indicated (YD►wt: *Mmp14*^*wt/wt*^ mice transplanted with BM from *Mmp14*^*Y573D/Y573D*^ mice; wt►YD: *Mmp14*^*Y573D/Y573D*^ mice transplanted with *Mmp14*^*wt/wt*^ BM). BM from non-transplanted *Mmp14*^*wt/wt*^ mice (wt) or *Mmp14*^*Y573D/Y573D*^ mice (YD) is shown as control. The PCR genotyping of non-transplanted mice with the three genotypes shown in the right panel is the same as shown in Figure 1 B, and is reported here to illustrate the expected pattern of BM with a mixed *Mmp14*^*wt/wt*^ and *Mmp14*^*Y573D/Y573D*^ genotype.

## TRANSPARENT METHODS

### Generation of MT1-MMP Y573D mice

To generate a mouse with the Y573D mutation in the MT1-MMP cytoplasmic tail we constructed a targeting vector as described in Figure 1 A. The construct was electroporated into W4 embryonic stem (ES) cells, followed by G418 and gancyclovir double selection. The W4 cells were provided to us by the Rodent Genetic Engineering of NYU School of Medicine, who carried out the blastocysts injections and subsequent procedures. Homologous recombination targeting events were identified in 8 ES cell colonies. Two of these, devoid of chromosome abnormalities by cytogenetic analysis, were used for injection into blastocysts of C57BL/6 mice (C57BL/6NTac; Taconic), and one gave germline transmission. Mice heterozygote for the mutation (*Mmp14*^*Y573/wt*^), identified by PCR analysis with primers flanking the *LoxP* sites (Fig. 1 A), were crossed with Rosa26Cre Deleter mice (C57BL/6NTac-*Gt(ROSA)26Sor*^*tm16(cre)Arte*^; Taconic) to excise the neo cassette. Homozygote *Mmp*^*Y573D/Y573D*^ mice were generated by heterozygote mating, and crossed back to wt C57BL/6 mice to remove the Cre recombinase gene. Sequencing of MT1-MMP cDNA from *Mmp*^*Y573D/Y573D*^ mouse tissue showed identity with wt MT1-MMP except for the Y573D mutation. Heterozygote mating produced wt, heterozygote and homozygote mice in the expected Mendelian ratios. The mice were genotyped by PCR using the same primers flanking the *loxP sites*, which afford identification of the three genotypes (Fig. 1 B). The sequences of these primers are:

Forward: GCT TGG CAG AGT GGA AAG AC
Reverse: GGG CAG TGA TGA AGG TGA GT

All the animal studies were carried out in strict accordance with the recommendations in the Guide for the Care and Use of Laboratory Animals of the National Institutes of Health. All procedures were approved by the Institutional Animal Care and Use Committee (IACUC) of New York University School of Medicine.

### Preparation of bone sections for histological analysis

Mice were euthanized by carbon dioxide narcosis, and the entire limb was removed by excising the femur at the joint in the upper extremity of the hip socket and the tibia at the ankle. The remaining skin was peeled off and the connective tissue removed. Limbs were placed in cold 4% paraformaldehyde for 24 h, and decalcified by incubation with Immunocal ^tm^ (Decal Corporation, Tallman, NY) or 10% EDTA for 72 h, with the decalcifying solution replaced with fresh one daily. Following decalcification, the limbs were rinsed 4 times for 30 min in PBS, 30 min in 0.85% NaCl and dehydrated for 30 min in 50% and 70% ethanol. Samples were embedded in paraffin blocks and sequentially cut into 5-μm sections, mounted onto slides and stained with the indicated reagents.

### Micro-computed tomography (CT)

Micro-CT was performed according to published guidelines (Bouxsein et al., 2010). Bones were scanned using a high-resolution SkyScan micro-CT system (SkyScan 1172, Kontich, Belgium). Images were acquired using a 10 MP digital detector, 10W energy (70kV and 142 mA), and a 0.5-mm aluminum filter with a 9.7-μm image voxel size. A fixed global threshold method was used based on the manufacturer’s recommendations and preliminary studies, which showed that mineral variation between groups was not high enough to warrant adaptive thresholds. The cortical region of interest was selected as the 2.0-mm mid-diaphyseal region directly below the third trochanter. The trabecular bone was assessed as the 2.0-mm region at the distal femur metaphysis; measurements included the bone volume relative to the total volume (BV/TV), bone mineral density (BMD), trabecular number (Tb.N), trabecular spacing (Tb.Sp), and trabecular thickness (Tb.Th).

### Immunohistochemistry

Paraffin-embedded specimens of brown adipose tissue (BAT) were cut and immunostained with antibody to uncoupling protein-1 UCP-1 (R&D Systems, cat. # MAB6158, Minneapolis, MN) at a concentration of 10 μg/mL by personnel of the Experimental Pathology Core of NYU School of Medicine. The antibody was validated using, as a negative control, sections of subcutaneous WAT devoid of BAT (Fig. 7 - figure supplement 2).

### Western blotting

Western blotting analysis was done as described (D’Alessio et al., 2008; Valacca, Tassone, & Mignatti, 2015) using the following antibodies and predetermined dilutions/concentrations. <u>MT1-MMP antibody</u>: Anti-MMP14 antibody [EP1264Y] rabbit mAb (Abcam, cat. # ab51074; Cambridge, MA), dilution: 1:5,000; <u>phospho-ERK1/2 antibody</u>: Phospho-p44/42 MAPK (Erk1/2) (Thr202/Tyr204) (197G2) Rabbit mAb (Cell Signaling Technology, cat. # 4377; Danvers, MA), dilution: 1:1,000; <u>phospho-Akt antibody</u>: Phospho-Akt (Ser473) (D9E) XP^®^ Rabbit mAb (Cell Signaling Technology, cat. # 4060), dilution: 1:2,000; <u>phospho-FAK antibodies</u>: Phospho-FAK (Tyr861) Recombinant Rabbit Monoclonal Antibody (26H16L4) (Invitrogen, cat. # 700154), concentration: 2 μg/ml; or Phospho-FAK (Tyr397) Polyclonal Antibody (Life technology, cat. # 44-624G), dilution: 1,1000; <u>total ERK1/2 antibody</u>: p44/42 MAPK (Erk1/2) (137F5) Rabbit mAb (Cell Signaling Technology, cat. # 4695), dilution: 1:1,000; <u>total Akt antibody</u>: Akt (pan) (C67E7) Rabbit mAb (Cell Signaling Technologies, cat. # 4691), dilution: 1:1,000; <u>total FAK antibody</u>: FAK Antibody C20 (Santa Cruz Biotechnology, cat. # sc-558), dilution: 1:200; <u>CD44 antibody</u>: mouse monoclonal antibody to mouse CD44 (DSHB; cat. # 5D2-27-s), dilution: 0.2 - 0.5 μg/ml; <u>ADAM9 antibody</u>: mouse anti-ADAM-9 monoclonal antibody (R&D Systems; cat. # MAB939); dilution: 1 μg /ml; <u>DDR1 antibody</u>: rabbit anti-DDR1 monoclonal antibody D1G6 (Cell Signaling Technology; cat. # 5583, dilution: 1:1,000; or Phospho-DDR1 (Tyr792) Antibody (Cell Signaling Technology; cat. #11994), dilution: 1:1,000; <u>α-tubulin antibody</u>: <u>rabbit anti-</u>tubulin monoclonal antibody 11H10 (Cell Signaling Technology; cat. #2125), dilution: 1:1,000; <u>β-tubulin antibody</u>: **β-Tubulin Antibody #2146** (Cell Signaling Technology; cat. #21465), dilution: 1:1,000; <u>lamin B antibody</u>: goat anti-lamin B antibody B-20 (Santa Cruz; cat. # sc-6217); dilution: 1:1,000; <u>rabbit IgG antibody</u>: anti-rabbit IgG, HRP-linked antibody (Jackson ImmunoResearch; cat. # AB_10015289), dilution 1:10,000, or (Cell Signaling Technologies, cat. # 7074), dilution: 1:10,000; <u>mouse IgG</u> antibody: anti-mouse HRP-linked antibody (Jackson ImmunoResearch; cat. # AB_2307391), dilution 1:10,000.

### Analysis of proMMP-2 activation by gelatin zymography

Confluent cells were incubated overnight with serum-free culture medium with or without addition of human recombinant MMP-2 (0.1 μg/ml; Anaspec, <u>Fremont, CA</u>). The conditioned medium was then treated with freshly prepared APMA solution (1 mM in DMSO; Sigma, St. Louis, MO) or DMSO as control, at 37° C degree for 1.5 h. Samples were analyzed as described [*Current Protocols in Immunology* 14.24.1-14.24.11, 2011] using precast SDS-polyacrilamide gels containing gelatin (Novex Zymogram Plus; ThermoFisher Scientific, Springfield, NJ).

### Collagen degradation assay

Single cell degradation of fibrillar collagen was analyzed as described (Sakr et al., 2018), using rat tendon or bovine collagen labeled with Alexa Fluor 488 (ThermoFisher Scientific). In a modification of this assay confluent cells were seeded onto Alexa Fuor 488 labeled fibrillar collagen in Dulbecco’s Modified Minimal Essential Medium (DMEM) supplemented with 10% fetal calf serum (FCS) to permit cell attachment. After 24 h incubation at 37° C, the cell layer was washed three times with serum-free DMEM and incubation was continued in serum-free phenol red-free to preclude MMP inhibition by serum inhibitors and interference with fluorescence measurement by phenol red. The medium was removed at 24 h intervals, immediately analyzed for Alexa Fluor 488 fluorescence with a fluorescence microplate reader, and replaced with the same volume of medium. Medium incubated with the labeled collagen in the absence of cells was used as a blanc whose fluorescence readings were subtracted from the samples’ readings at each time point. The time of the first addition of serum-, phenol red-free DMEM was considered as time 0.

### Cell migration and collagen invasion assays

For migration and invasion assays the polycarbonate porous membranes of inserts of 24-well Transwell plates (Nunc, ThermoFisher Scientific, 8-μm pores, 0.47 cm^2^ culture area) were coated with either 50 ul of either PBS, for migration assays, or 0.1% calf skin collagen (Sigma-Aldrich) for invasion assays. After incubation at 37° C for 45 min and washing three times with serum-free DMEM, 1.4 - 1.6 × 104 cells (corresponding to confluent cultures) the membranes were seeded into each insert in 100 ul of DMEM supplemented with 10% FCS. Six hundred microliters of the same medium was added to the bottom of each well, and the plates were incubated at 37° C for 2 h to allow cell attachment. The inserts were gently rinsed three times to remove the serum, and 100 ul of serum-free DMEM was added into each insert, so as to generate a gradient of 0-10% serum across the porous membranes. Following incubation at 37° C for 24 h, the cultures were fixed with 4% paraformaldehyde for 15 min and stained with 10% Giemsa, after which the cell layer on top of the membranes was removed by wiping with a Q-tip. The membranes were detached from the inserts, and mounted upside-down onto microscope slides. The migrated cells were counted under a microscope with 20X magnification in 10 random fields per membrane. Triplicate samples were used.

### Flow cytometry

Flow cytometric analysis of MT1-MMP was performed exactly as we described (Harbi et al., 2016) using MMP14 Alexa Fluor® 488-conjugated antibody (R&D systems, cat. # FAB9181G) or mouse IgG2B Alexa Fluor® 488-conjugated isotype control (R&D systems, cat. #IC0041G) following the manufacturer’s instructions.

### Isolation, culture and differentiation of BM SSC

Cultures of BM SSC were established by a modification of the method described (Soleimani & Nadri, 2009). BM was flushed from cut femurs and tibiae using 3-ml syringes with Minimum Essential Medium Alpha (αMEM without ribonucleosides or deoxyribonucleosides; Gibco, Life Technologies cat. # 12561-056, Grand Island, NY) supplemented with 15% fetal calf serum (FBS; Atlanta Biologicals, cat. # S11150, Flowery Branch, GA) and antibiotics (Penicillin-Streptomycin). After separating the BM into a single-cell suspension, the cells were passed through a 70-μm cell strainer, incubated for 2 min in NH4Cl red blood cell lysis solution (StemCell Technologies, cat. # 07800, Vancouver, Canada), and seeded into culture plates in αMEM supplemented with 15% FBS and antibiotics. Three hours later non-adherent cells were removed by replacing the medium with fresh complete medium. This procedure was repeated twice a day for the first 72 h. Subsequently, the adherent cells were washed with phosphate buffer saline (PBS) and incubated with fresh medium every 3-4 days until the cultures became subconfluent. To induce differentiation the cells were seeded into 24-well plates (5 × 10^4^ cells/well). After 24 h, the culture medium was substituted with osteoblast or adipocyte differentiation medium, and incubation was continued for the indicated times, changing the medium with freshly prepared medium twice a week. For chondrocyte differentiation the cells were seeded into 48-well plates as a pellet (2.5 × 10^5^ cells/well). The medium was substituted with freshly prepared differentiation medium 2 h after seeding, and subsequently three times a week. Differentiation medium consisted of αMEM supplemented with 10% FBS, penicillin/streptomycin and, for osteoblast differentiation: dexamethasone 1 μM (Millipore Sigma, cat. # D4209; St. Louis, MO), beta-glycerolphosphate 20 mM (Millipore Sigma, cat. # 500200), L-ascorbic acid 50 μM (Millipore Sigma, cat. # A5960); for adipocyte differentiation: dexamethasone 1 μM, indomethacin 50 μM (Millipore Sigma, cat. # I7378), 3-Isobutyl-1-methylxanthine (IBMX) 500 nM (Millipore Sigma, cat. # I5879), insulin 5 μg/ml (Millipore Sigma, cat. # I9278); and for chondrocyte differentiation: sodium pyruvate 100 μg/ml, ascorbic acid 50 μg/ml, dexamethasone 0.1 μM, insulin, transferrin, selenium (ITS) mix 1X (Millipore Sigma, cat. # I3146) and bone morphogenetic protein-2 (BMP2) 300 ng/ml (Peprotech, cat. # 120-02-2UG, Rocky Hill, NJ). The SSC used for differentiation experiments were no older than passage 3 in culture.

### CH310T1/2 cell culture, transfection and differentiation

C3H10T1/2 cells obtained from the American Type Culture Collection (ATCC CLL226, Gaithersburg, MD) were grown in Dulbecco’s MEM (DMEM; Gibco, Life Technologies, cat. # 10566-016) supplemented with 10% FCS (Atlanta Biologicals). Cells at 50-70% confluency were transfected by overnight incubation with the indicated cDNAs in complete growth medium using TransIT®-LT1 - Transfection Reagent according to the manufacturer’s instructions (Mirus Bio, LLC, cat. # MIR2300; Madison, WI). Stable transfectants were selected in medium containing 500 μg/ml hygromycin B (InvivoGen, cat. # ant-hg-1; San Diego, CA). Pools of resistant cell colonies were characterized for MT1-MMP expression by reverse transcription-PCR and Western blotting, and used for differentiation assays. To induce differentiation the cells were seeded into 24-well plates at a density of 2.5 × 10^4^ cells/well and then shifted to DMEM with low glucose (1 g/L; Gibco, Life Technologies, cat. # 11885076) with the same FCS concentration and supplements used for BM-derived SSC differentiation.

### Quantitative PCR analysis of gene expression in differentiating cells

Cells were lysed in Trizol Reagent (Invitrogen, Waltham, MA) and total RNA was isolated with RNeasy (Qiagen, Valencia, CA), according to the manufacturer’s instructions. RNA was quantified using a Nanodrop 2000 spectrophotometer (ThermoFisher Scientific, Waltham, MA), and 1 μg of total RNA was used for cDNA preparation using SuperScript III (Invitrogen, Life Technologies). Quantitative real-time PCR reactions were performed with SYBR green PCR reagents using the ABI Prism 7300 sequence detection system (Applied Biosystems-Life Technologies, Waltham, MA). Relative gene expression levels were calculated using the 2 delta Ct method (Livak & Schmittgen, 2001). Target mRNA levels were normalized to the geomean GAPDH and HPRT1.

### Primers for qPCR analysis of differentiation markers: Osteoblast

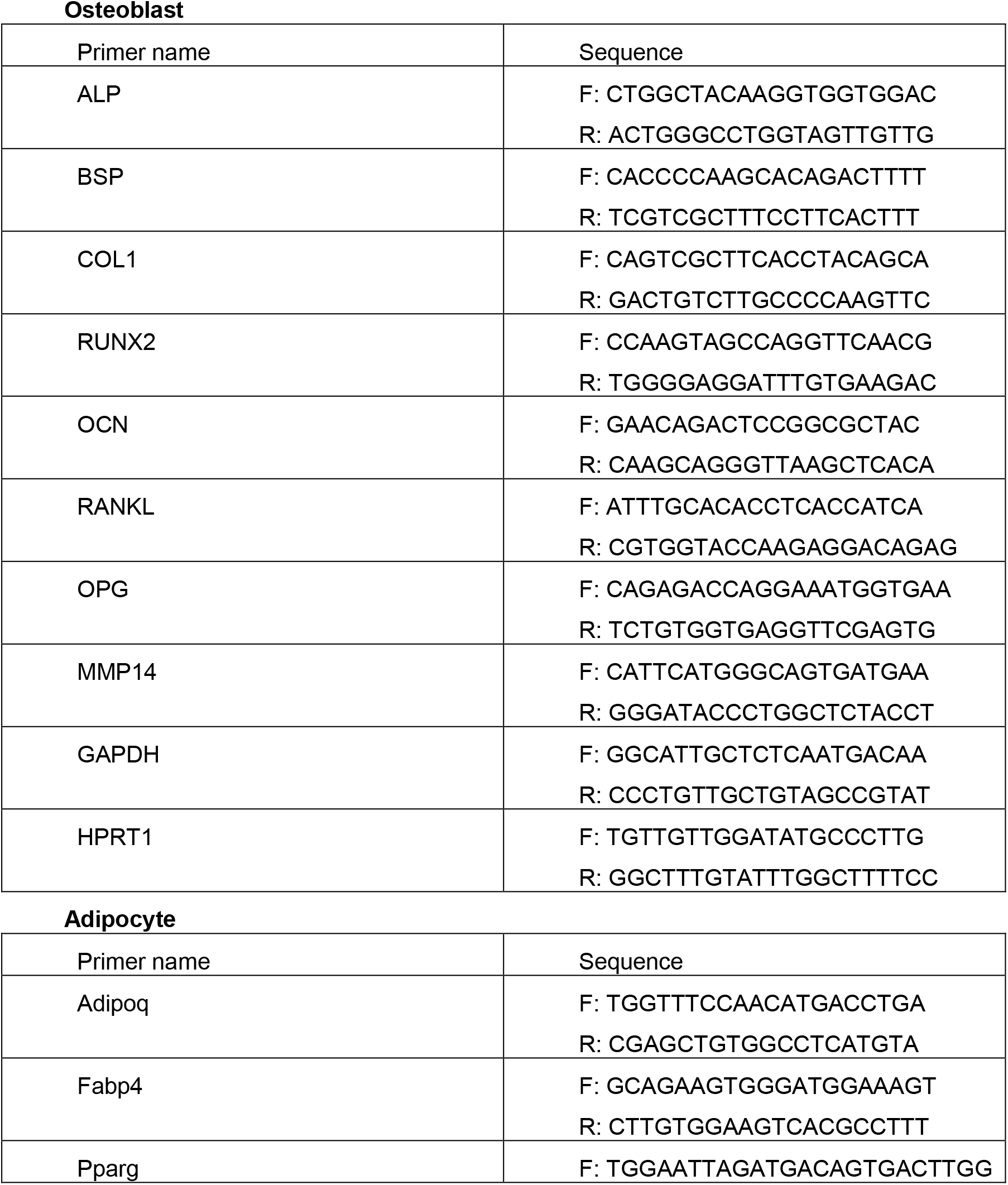
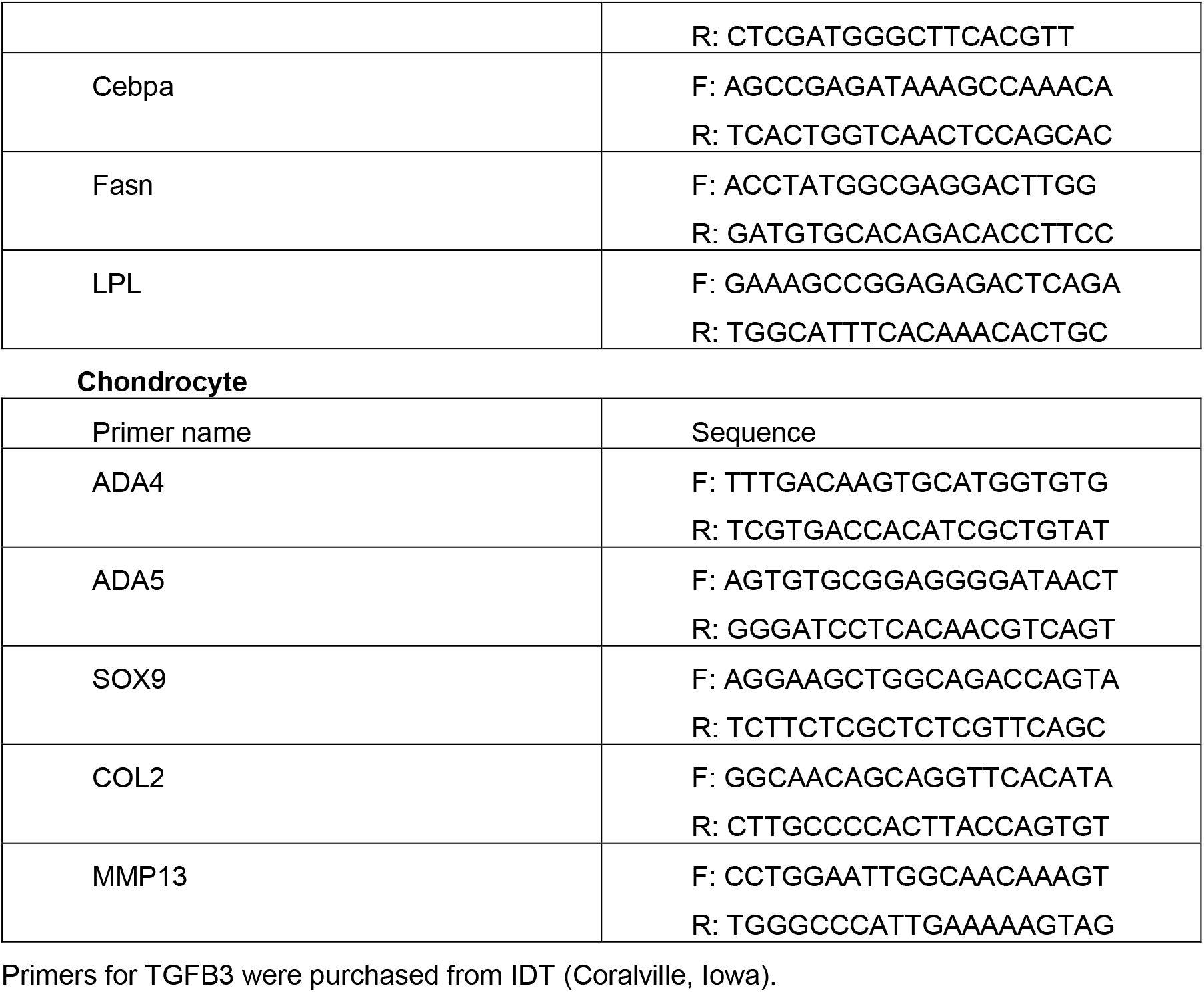

### Analysis of circulating hormones and cytokines

Serum and/or plasma levels of the indicated hormones and cytokines were measured by multiplex ELISA kits (Millipore, Burlington, MA) at the High-Throughput Biology Laboratory of NYU School of Medicine.

### RNA-Seq and data analysis

The indicated tissues and SSC were isolated from *Mmp14*^*wt/wt*^ and *Mmp14*^*Y573D/Y573*^ mice (3 mice/tissue/genotype) and total RNA was extracted using the RNeasy kit (Qiagen). The subsequent steps were carried out by personnel of the Genome Technology Center of NYU School of Medicine. RNA-Seq libraries were prepared with the TruSeq sample preparation kit (Illumina, San Diego, CA). Sequencing reads were mapped to the mouse genome using the STAR aligner (v2.5.0c) (Dobin et al., 2013). Alignments were guided by a Gene Transfer Format file (Ensembl GTF version GRCh37.70). The mean read insert sizes and their standard deviations were calculated using Picard tools (v.1.126). Read count tables were generated using HTSeq (v0.6.0) (Anders, Pyl, & Huber, 2015), normalized to their library size factors with DESeq (v3.7) (Anders & Huber, 2010), and differential expression analysis was performed. Read Per Million (RPM) normalized BigWig files were generated using BEDTools (v2.17.0) (Quinlan & Hall, 2010) and bedGraphToBigWig tool (v4). Statistical analyses were performed with R (v3.1.1), GO analysis with David Bioinformatics Resources 6.8 (Huang da, Sherman, & Lempicki, 2009a, 2009b) and GSEA with the hallmarks (h.all.v6.1.symbols.gmt) gene sets of the Molecular Signature Database (MSigDB) v6.1 (Mootha et al., 2003; Subramanian et al., 2005).

### BM transplant

Mice of 3 and 5 months of age were used for BM transplant experiments. The BM of female mice was transplanted into age-matched male mice. The recipient mice were lethally irradiated with a single dose of 9 Gy. Twenty-four hours later BM cells (BMCs) were collected from the tibias and femurs of donor mice, suspended in PBS containing 2% heat-inactivated FBS, and 5 × 10^6^ BMCs in 200 μl were injected into the recipient mice via the tail vein. Eight weeks later the mice were subjected to DEXA scanning analysis and then sacrificed. To verify BM engraftment we transplanted female mice with male BM, in order to characterize the recipients’ BM by immunohistochemistry and/or flow cytometry with antibody to UTY, a Y chromosome marker (Wang et al., 2013). However, several commercially available antibodies failed to recognize UTY in murine BM cells, probably because this marker is not expressed in these cells. For the same reasons we were not able to identify the genotype of transplanted animals’ WAT or BAT by immunohistochemistry with UTY antibody, or RNA *in situ* hybridization. Therefore, we genotyped the BM of recipient mice for the *Mmp14*^*Y573D/Y573D*^ mutation using the PCR primers designed for genotyping our mice. We reasoned that partial engraftment of donor’s BM would result in both the *Mmp14*^*wt/wt*^ and *Mmp14*^*Y573D/Y573D*^ genotypes, with a pattern of PCR products identical to that of *Mmp14*^*Y573D/wt*^ mice (Fig. 1 B). In addition, running the PCR reaction for a high number of cycles would afford detection of small residual amounts of BM from recipient mice. By this method, the results (Fig. 10 - figure supplement 1) showed virtually complete replacement of the recipient’s BM with the donors’ BM. However, we could not use this method to identify donor-derived adipocytes in the transplanted animals’ WAT or BAT as these tissues contain circulating cells - e.g. macrophages - from the BM.

### Statistical analysis

For the BM transplant experiments, to calculate the number of mice required in each group to achieve statistical significance for the expected differences between controls and experimental groups, we used the analysis of Power of Test based on the variance observed in our previous analyses of our mice. In our characterization of the bone, cartilage and fat phenotypes we found differences between MT1-MMP Y573D wt mice ranging 25% - 75% in the mean (m) values of the various parameters, with standard deviations (SD) ranging 25% - 50%. Therefore, we assumed m = 50% and SD = 35%. To calculate the number of animals needed we used the formula: n = (Za +Zb)^2^ × 2 (%SD)^2^/ (%D)^2^, where: n is number of animals per group, (Za + Zb)^2^ for a 2-Tailed test with power of 90% at p=0.05 is 10.5, %SD is the percent standard deviation, and %D is the assumed percent difference between control and experimental group(s). Thus, n = 10.5 × 2(35)^2^ / 50^2^ = 10.29. Therefore, we planned to use 11 mice for each experimental and control group. The statistical analysis of the results of all the experiments was performed with the two-tailed Student’s t-test (unless otherwise indicated) using GraphPad Prism, Version 7.05.

## Notes

### Competing Interest Statement

The authors have declared no competing interest.

### Summary of Updates

Membrane-type 1 matrix metalloproteinase (MT1-MMP, MMP-14), a transmembrane proteinase with a short cytoplasmic tail, is a major effector of extracellular matrix (ECM) remodeling. Genetic silencing of MT1-MMP in mouse (Mmp14-/-) and man causes dwarfism, osteopenia, arthritis and lipodystrophy, abnormalities ascribed to defective collagen turnover. We have previously shown non-proteolytic functions of MT1-MMP mediated by its cytoplasmic tail, where the unique tyrosine (Y573) controls intracellular signaling. The Y573D mutation blocks TIMP2/MT1-MMP-induced Erk1/2 and Akt signaling without affecting proteolytic activity. Here we report that a mouse with the MT1-MMP Y573D mutation (Mmp14Y573D/Y573D) shows abnormalities similar to, but also different from those of Mmp14-/- mice. Skeletal stem cells (SSC) of Mmp14Y573D/Y573D mice show defective differentiation consistent with the mouse phenotype, which is rescued by wild-type SSC transplant. These results provide the first in vivo demonstration that MT1-MMP modulates bone, cartilage and fat homeostasis by controlling SSC differentiation through a mechanism independent of proteolysis.

